# Spindle reorientation in response to mechanical stress is an emergent property of the spindle positioning mechanisms

**DOI:** 10.1101/2022.02.13.480269

**Authors:** Manasi Kelkar, Pierre Bohec, Matthew Smith, Varun Sreenivasan, Ana Lisica, Leo Valon, Emma Ferber, Buzz Baum, Guillaume Salbreux, Guillaume Charras

**Affiliations:** London Centre for Nanotechnology, University College London, London, WC1H 0AH, UK; The Francis Crick Institute, 1 Midland Road, London NW1 1A, UK; Centre for Developmental Neurobiology, Institute of Psychiatry, Psychology and Neuroscience, King’s College London, London SE1 1UL, UK; MRC Centre for Neurodevelopmental Disorders, King’s College London, London SE1 1UL, UK; Institut Pasteur, 25-28 rue du Dr Roux, 75015 Paris, France; MRC Laboratory for Molecular Cell Biology, University College London, Gower Street, London WC1E 6BT, UK; Division of Cell Biology, MRC Laboratory of Molecular Biology, Cambridge UK; Institute for the Physics of Living Systems, University College London, London, WC1E 6BT, UK; Department of Genetics and Evolution, Geneva, Switzerland; Department of Cell and Developmental Biology, University College London, London, WC1E 6BT

## Abstract

Proper orientation of the mitotic spindle plays a crucial role in embryos, during tissue development, and in adults, where it functions to dissipate mechanical stress to maintain tissue integrity and homeostasis. While mitotic spindles have been shown to reorient in response to external mechanical stresses, the subcellular cues that mediate spindle reorientation remain unclear. Here, we have used a combination of optogenetics and computational modelling to better understand how mitotic spindles respond to inhomogeneous tension within the actomyosin cortex. Strikingly, we find that the optogenetic activation of RhoA only influences spindle orientation when it is induced at both poles of the cell. Under these conditions, the sudden local increase in cortical tension induced by RhoA activation reduces pulling forces exerted by cortical regulators on astral microtubules. This leads to a perturbation of the torque balance exerted on the spindle, which causes it to rotate. Thus, spindle rotation in response to mechanical stress is an emergent phenomenon arising from the interaction between the spindle positioning machinery and the cell cortex.

## Introduction

The orientation of the mitotic spindle determines the axis of cell division **[1, 2]**. Cell division orientation must be carefully regulated during embryonic development to dictate proper cell fate as well as in adult organisms to maintain tissue homeostasis or enable adaptation to environmental changes **[3–6]**. In tissues, division orientation has been shown to homogenise cell packing **[7]** and to prevent the build-up of excessive mechanical stress, which may endanger tissue integrity. In monolayered epithelia, cell divisions parallel to the substratum are essential to maintain and direct in-plane tissue growth **[8, 9]**. When divisions have no favoured orientation, growth is isotropic; whereas orientation along one specific axis can lead to tissue elongation **[10, 11]**. As a consequence, the cues and mechanisms regulating the orientation of cell division have been the focus of much research.

Mitotic spindles in both isolated cells and tissues can alter their orientation in response to a variety of cues. Depending on the experimental conditions, spindle orientation can be influenced by cell shape, junctional cues, signals from the extracellular matrix, the stress field, or a combination of these cues **[6, 12–21]**. To probe the cellular-scale response to tissue deformation, studies *in vivo* and *in vitro* have applied uniaxial deformation to isolated cells as well as epithelia and shown cell division to be oriented along the stretch axis **[6, 17, 22, 23]**. Outside of a tissue context, elongated metaphase cells that are subjected to a uniaxial stretch perpendicular to their long axis reorient their spindle to a position intermediate between the directions provided by the shape and stress cues **[17]**. In developing tissues, such as the zebrafish enveloping layer, a stress field applied perpendicular to the spindle axis causes reorientation **[14]**. Collectively, these studies indicate that spindles can respond to externally applied deformation and stresses. However, how tissue-scale deformation impacts the tension distribution in the cortex of a mitotic cell is unclear. Moreover, whether and how spindles are able to respond to tension inhomogeneities within the actin cortex remains unknown. Indeed, the reorientation of spindles could arise from the activation of mechanosensitive signalling pathways or could emerge from changes in the balance of forces exerted by the spindle positioning apparatus.

To address the role of the mitotic cortex in spindle orientation, we use optogenetic activators of contractility to generate inhomogeneous tension within the cortex of mitotic cells and investigate the spindle response. Remarkably, an increase in cortical myosin and tension specifically at cell poles results in reorientation of the metaphase spindle away from these regions, subsequently affecting the orientation of cell division. Spindle rotation arises due to a local reduction of the pulling forces exerted by cortical regulators on astral microtubules. Experiments and mathematical modelling suggest that rotation emerges from the interaction between the spindle positioning machinery and the cell cortex. Our data therefore suggest that mitotic spindles can respond to inhomogeneities in myosin activity and cortical tension to orient division away from regions of high tension.

## Results

### Increase in cortical RhoA activity increases cortical tension

The tension distribution that arises in the cortex of mitotic cells in response to the application of a stress field remains poorly understood. However, application of uniaxial stress to rounded mitotic cells likely leads to inhomogeneous tension in the cortex. To investigate the response of mitotic spindles to such tension inhomogeneities, we modulated cortical tension in mitotic cells by regulating the activity of RhoA at the plasma membrane using optogenetics. To this end, we used a previously established light-gated CRY2/CIBN dimerisation system **[24, 25]**. In this actuator, the DH-PH domain of a RhoA specific GEF, p115-RhoGEF/Arhgef1 **[26]**, is fused to CRY2-mCherry and stably expressed in MDCK cells alongside CIBN-GFP which is targeted to the plasma membrane with a CAAX domain (Fig. 1A). Exposure to blue light induces a conformational change in CRY2 causing it to bind CIBN and re-localise the DH-PH domain to the plasma membrane (Fig. 1A). With this system, recruitment to the membrane can be restricted to subcellular regions as small as 5 μm **[27]** and optogenetic re-localisation of GEF DH-PH to the membrane increases RhoA activity **[25]**. We reasoned that RhoA activation should lead to an increase in myosin contractility and subsequently an increase in cortical tension. To verify this, we measured cortical tension in rounded cells before and after exposure of the whole cell to blue light using atomic force microscopy (AFM) (Fig. S1A). These measurements revealed a two-fold increase in tension following optogenetic activation (Fig. 1B and C) confirming that GEF DH-PH re-localisation modulates cortical tension.

**Figure1:**
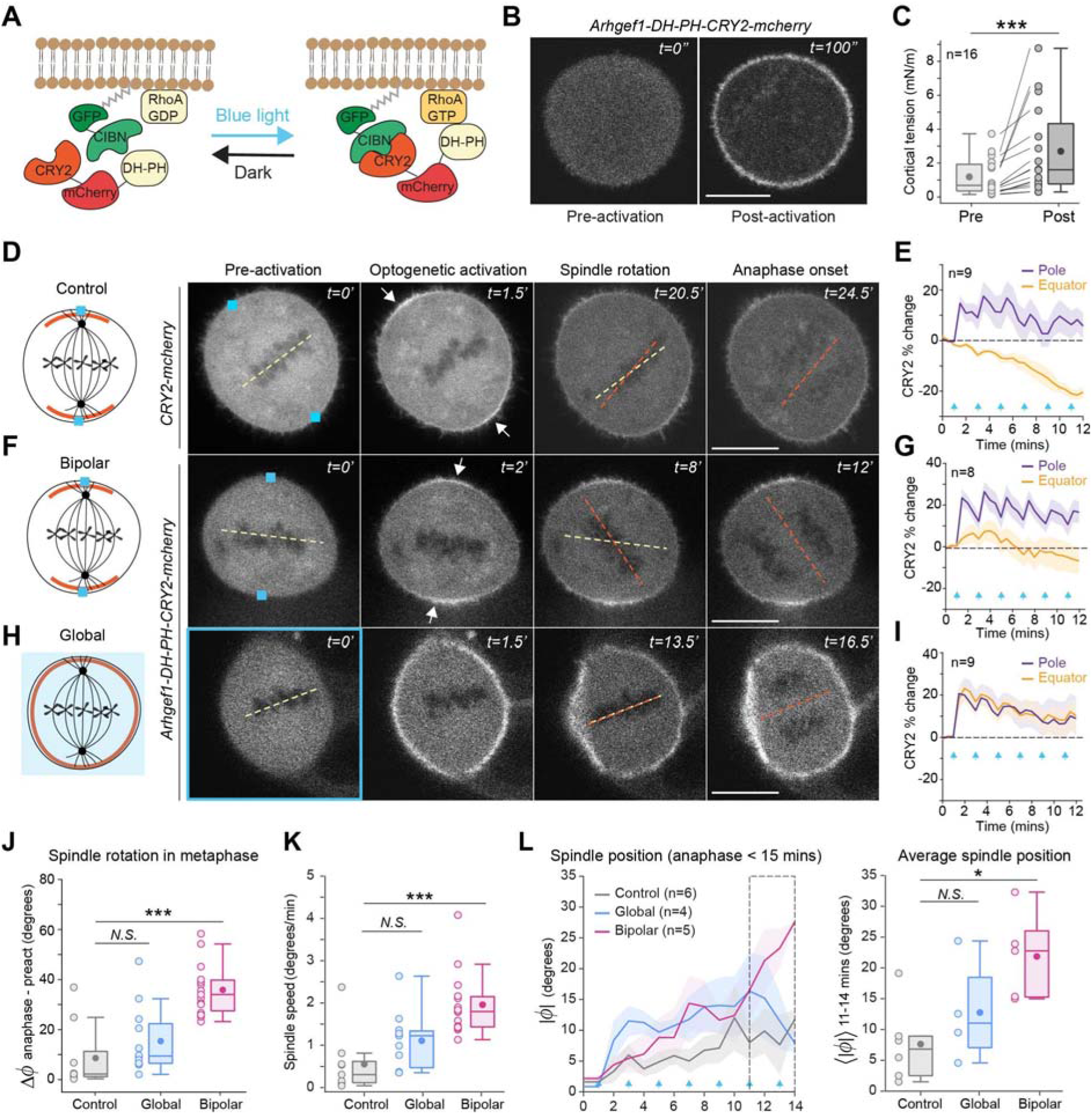
Mitotic spindle orientation responds to inhomogeneities in RhoA activity and cortical tension. In all images, scale bars = 10μm. Boxplots show the median (dark line), interquartile range, mean (dark filled circle) and individual data points (light filled circle). A. Schematic depicting the light-gated CRY2/CIBN dimerisation system used to regulate RhoA activity. The N-terminal of CIB (CIBN) is fused to GFP and targeted to the plasma membrane using a CAAX motif. CRY2 is cytosolic, tagged with mCherry and fused to the DH-PH domain of a RhoA-specific GEF-Arhgef1/p115Rho-GEF. Upon illumination with 473nm blue light, CRY2 undergoes a conformational change and binds to CIBN, thereby relocating the DH-PH domain of the GEF to the plasma membrane where it can activate RhoA. After illumination is stopped, CRY2 undergoes a slow detachment from CIBN and returns to the cytosol. B. Representative confocal images of rounded interphase MDCK cells co-expressing CIBN-GFP-CAAX and Arhgef1-GEF-DH-PH-CRY2-mCherry viewed in the mCherry channel at t=0 (preactivation) and t=100sec after blue light illumination (post-activation). Time is in seconds. C. Boxplot showing cortical tension in cells as in (B) measured by atomic force microscopy pre- and post-exposure to blue light (n=16 cells). Wilcoxon sign-rank test: *p*= 4.7×10^-4^ (***). D. Left: Schematic showing the localisation of CRY2-mCherry (Control), (red), after bipolar optogenetic activation in metaphase cells. Right: Time series showing localisation of CRY2-mCherry (control) in cells before and after bipolar optogenetic activation until anaphase onset. Yellow and orange dashed lines indicate the position of the DNA in the metaphase plate before activation and at anaphase onset. Blue squares indicate the region of activation, white arrows point to regions of CRY2-mCherry accumulation. Time is in minutes. E. Temporal plot showing the percentage change in CRY2-mCherry (control) intensity at the poles and equator after bipolar activation. Plot shows mean ± SEM (n=9 cells). Blue arrows indicate the times of blue light stimulation. F. Left: Schematic showing the localisation of Arhgef1-DH-PH-CRY2-mCherry (red) after bipolar optogenetic activation in metaphase cells. Right: Time series showing localisation of GEF-DH-PH-CRY2-mCherry in cells before and after bipolar optogenetic activation until anaphase onset. Yellow and orange dashed lines indicate the position of the DNA in the metaphase plate before activation and at anaphase onset. Blue squares indicate the region of activation, white arrows point to regions of GEF-DH-PH-CRY2 accumulation. Time is in minutes. G. Temporal plot showing the percentage change in Arhgef1-DH-PH-CRY2-mCherry intensity at the poles and equator after bipolar activation. Plot shows mean ± SEM (n=8 cells). Blue arrows indicate the times of blue light stimulation. H. Left: Schematic showing the localisation of Arhgef1-DH-PH-CRY2-mCherry (red) after global optogenetic activation in metaphase cells. Right: Time series showing localisation of GEF-DH-PH-CRY2-mCherry in cells before and after global activation until anaphase onset. Time is in minutes. I. Temporal plot as in (G) after global optogenetic activation (n=9 cells). J. Boxplot showing spindle rotation in metaphase for cells after control (n=9), global (n=12) and bipolar (n=15) activation. Kruskal-Wallis and post-hoc Dunn’s test: Control vs Bipolar: *p*<0.001 (***); Control vs Global: *p*=0.4168 (N.S). K. Boxplot showing spindle rotation speed after activation until anaphase onset for cells after control (n=9), global (n=11) and bipolar (n=15) activation. ANOVA and Tukey-Kramer test: Control vs Bipolar: *p*=0.0002 (***); Control vs Global: *p*=0.21 (N.S). L. Left: Trajectories of absolute spindle angles before and for 14 minutes after optogenetic activation for a subset of cells, whose anaphase occurs within 15 minutes of activation. Right: Boxplot showing an average of the absolute spindle angle between 11-14 minutes. Control (n=6), bipolar (n=5), global (n=4). ANOVA and Tukey-Kramer test: Control vs Bipolar: *p*=0.017 (*); Control vs Global: *p*=0.527 (N.S).

Next, we determined if local accumulation of GEF DH-PH translated into a local increase in cortical tension. Cell shape is controlled by cortical tension, which depends strongly on myosin contractility **[28]**. Therefore, we reasoned that changes in cell shape in response to localised optogenetic activation should indicate a change in cortical tension. To this end, we illuminated specific regions in MDCK cells synchronised in metaphase with brief pulses of a 473-nm blue laser every 2 minutes to maintain a constant level of CRY2, and therefore GEF DH-PH, at the membrane. Optogenetic re-localisation of DH-PH to both poles led to a decrease in the length of the polar axis and an increase in the length of the equatorial axis causing a reduction in the ratio of polar/equatorial axis (Fig. S1B and C) and flattening of the cell at the poles. Along with global AFM measurements showing an increase in cortical tension in response to global activation (Fig. 1B and C), these data indicate that localised optogenetic activation locally increases cortical tension.

### Mitotic spindle orientation responds to local differences in cortical tension induced by RhoA activation

To investigate the response of mitotic spindles to local inhomogeneities in cortical tension, we targeted our optogenetic actuator either to cell poles alone, the equator alone or isotropically across the whole cell and examined the spindle response until the onset of cytokinesis. Localised activation in small square regions (0.62μm x 0.62μm) specifically at cell poles (bipolar) led to ~20% increase in GEF accumulation in those regions without significantly affecting GEF levels at the equator (Fig. 1F and G; Fig. S2F). Remarkably, this resulted in an average ~35° rotation of the mitotic spindle away from its original orientation, subsequently causing a change in the division axis in these cells (Fig. 1F and J). Spindle reorientation was specific to GEF re-localisation, as it was not observed in cells expressing CRY2-mCherry without the DH-PH domain (control) (Fig. 1D, E and J; Fig. S2C). Following this observation, we asked if any spatial heterogeneity or temporal change in RhoA activity and cortical tension was sufficient to cause spindle reorientation. Although localised optogenetic activation at the equator (equatorial) (Fig. S2A) or whole cell optogenetic activation (global) (Fig. 1H) both led to GEF accumulation in those regions (Fig. S2B, D, E, and Fig. 1I), neither perturbed the average spindle orientation nor the division axis (Fig. 1J; Fig. S2G). Furthermore, spindle rotation only occurred in metaphase (Fig. 1J) with very little change in orientation from anaphase until cytokinesis (Fig. S2H), suggesting that this phenomenon may be specific to metaphase. Consistent with this, when we carried out optogenetic activation at the poles in early anaphase, no spindle rotation was observed, indicating that rotation was no longer possible after anaphase onset (Fig. S2J). Thus, our data show that the spindles specifically respond to a local increase in RhoA activity and cortical tension at cell poles during metaphase.

Following these observations, we sought to further characterise the spindle response to optogenetic activation in metaphase. In theory, the final angle that a spindle reaches should be influenced by its rotation speed and/or the time available for rotation until anaphase onset. The former proved to be the case as the speed of spindle rotation, measured from optogenetic activation until anaphase onset, was significantly higher in cells subjected to bipolar activation than under control condition (Fig. 1K). Moreover, optogenetic activation of the GEF at cell poles did not lead to a significant difference in the time until anaphase onset (Fig. S2I). Finally, when we measured the temporal evolution of spindle position by plotting the spindle angle trajectories averaged over all cells up to 14 minutes post-activation, we did not identify a significant difference in the final spindle position between control and bipolar activation conditions (Fig. S2K). As the time until anaphase onset upon bipolar GEF activation was more variable than in control cells (Fig. S2I), we decided to analyse instead the temporal evolution of spindle position in a subset of cells that enter anaphase within 15 minutes of optogenetic activation. Again in this case, spindles underwent a greater rotation following bipolar GEF activation than under control condition (Fig. 1L).

Taken together, these results demonstrate that the increased levels of RhoA activity and cortical tension induced by the local recruitment of GEF DH-PH to cell poles increase the rotation speed of the spindle to reorient divisions.

### Mitotic spindles rotate away from myosin-enriched, tensed cortical regions

Next, we sought to understand the molecular changes in the cortex leading to spindle rotation. We first verified that localised optogenetic actuation of GEF DH-PH indeed led to local activation of RhoA by imaging the RhoA biosensor iRFP-AHDPH **[29]**. Bipolar activation led to a ~15% increase in RhoA biosensor intensity at the poles without significantly affecting equatorial levels (Fig. S3A). RhoA activates Rho-associated coiled-coiled kinase (ROCK) **[30]**, which increases myosin activity through two pathways, phosphorylating and activating the regulatory light chain of myosin-II while simultaneously inhibiting myosin phosphatase **[31, 32]**. RhoA also acts on the F-actin scaffold directly by activation of the cortical actin nucleator mDia1 **[33, 34]** and indirectly via ROCK, which has been shown to inactivate cofilin via phosphorylation through LIMK **[35]**. Thus, a change in RhoA activity can potentially regulate cortical tension by activating myosins and/or by changing the F-actin scaffold.

To determine how RhoA increased cortical tension, we examined the response of cortical myosin and F-actin to optogenetic actuation of GEF DH-PH. Bipolar activation led to an increase in MRLC-iRFP intensity at cell poles, which reached a steady state within ~3 minutes of activation (2.8 ±0.74 min, n=8 cells), marking it as a potential early event that preceded spindle rotation away from the poles (Fig. 2A). Consistent with GEF re-localisation (Fig. 1G), we observed a significant increase in myosin intensity of ~15% at the poles whereas no significant changes were observed in the future pole or at the equator (Fig. 2B, C and D). In contrast, F-actin levels imaged using LifeAct-iRFP displayed no clear changes in intensity (Fig. S3B). Thus, these data indicate that local activation of RhoA leads to local differences in myosin levels in the cortex that underlie inhomogeneities in cortical tension.

**Figure2:**
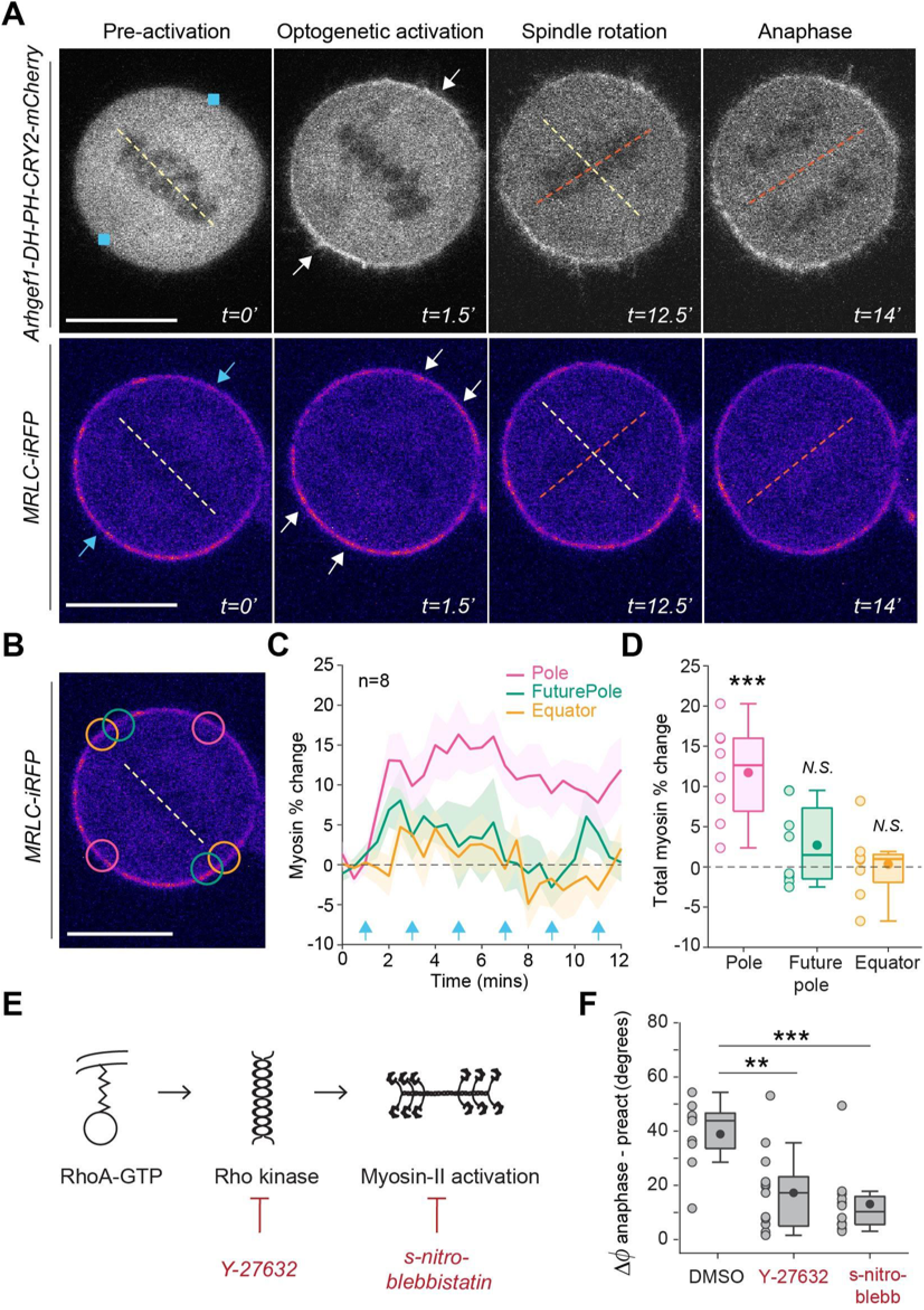
Mitotic spindles rotate away from myosin-enriched tensed cortical regions. A. Top panel: Time series showing localisation of Arhgef1-DH-PH-CRY2-mCherry in cells before and after bipolar optogenetic activation until anaphase onset. Bottom panel: Time series showing the corresponding localisation of MRLC-iRFP with low intensities appearing in black and high intensities in magenta. Yellow and orange dashed lines indicate the position of the DNA in the metaphase plate before activation and at anaphase onset respectively. Blue squares (top panel) and blue arrows (bottom panel) indicate the region of activation, white arrows point to regions of GEF-DH-PH accumulation (top panel) and myosin accumulation (bottom panel). Scale bar = 10μm. Time is in minutes. B. MRLC-iRFP image at t=0 as in (A-bottom panel) showing the regions used to measure fluorescence intensity in (C-D), with pole (pink), future pole (green) and equator (yellow). Scale bar is 10μm. C. Temporal plot showing the percentage change in MRLC-iRFP intensity at the pole, future pole and equator after bipolar activation. Plot shows mean ± SEM (n=8 cells). Blue arrows indicate the times of blue light stimulation. D. Boxplot showing the total percentage change in MRLC-iRFP intensity at the pole, future pole and equator until 11 minutes post-activation for cells as in (C). Boxplot shows median (dark line), interquartile range, mean (dark filled circle) and individual data points (light filled circle), (n=8 cells). Student *t*-test against 0% change: Pole: *p*=1.5×10^-4^ (***), Future pole: *p*=0.0840 (N.S), Equator *p*=0.8247 (N.S). E. Schematic depicting part of the signalling downstream of RhoA. Active RhoA-GTP activates Rho-kinase that, in turn, activates myosin-II by direct phosphorylation and inhibition of myosin phosphatase. Y-27632 and s-nitro-blebbistatin block myosin contractility by inhibiting Rho-kinase and myosin ATPase activity, respectively. F. Boxplot showing spindle rotation in metaphase for control cells treated with DMSO (n=9), 50μM Y-27632 (n=13) and 20μM s-nitro-blebbistatin (n=12) respectively. Boxplot shows median (dark line), interquartile range, mean (dark filled circle) and individual data points (light filled circle). ANOVA and Tukey-Kramer test: DMSO vs s-nitro-blebbistatin: *p*= 0.0005 (***); DMSO vs Y-27632: *p*= 0.0028 (**).

To determine the role of myosin generated cortical tension in spindle rotation, we performed optogenetic activation experiments in the presence of inhibitors of myosin activity **(Fig2E)**. Inactivation of Rho-kinase using Y-27632 or direct inhibition of myosin activity using photostable s-nitro-blebbistatin **[36]**, both abolished spindle rotation, signifying that myosin activity at the poles is essential to enable spindle rotation (Fig. 2F). As both treatments decrease cortical tension **[37, 38]**, spindles may therefore either be sensitive to myosin activity directly or to the increase in cortical tension that it causes.

Taken together, our results suggest that localised RhoA activation at the poles acts through a local increase in myosin enrichment and cortical tension to induce spindle rotation.

### Spindle rotation depends on cortical localisation of NuMA and pulling forces exerted by Dynein

We next investigated the role of the previously identified spindle positioning machinery in driving rotation. Mitotic spindle positioning depends on a conserved complex of Gαi, LGN and NuMA localised at the plasma membrane that recruits the microtubule minus-end directed motor protein Dynein **[8, 39, 40]** (Fig. 3A). Astral microtubules that extend from the spindle poles to the cortex play a key role in spindle positioning by exerting pushing forces on the cortex generated by microtubule growth and pulling forces that are associated with a combination of microtubule shrinkage and Dynein motor activity **[41–44]**. To investigate the contribution of these proteins to spindle rotation, we performed inactivation experiments to block each component in turn. Inhibition of Dynein activity using CiliobrevinD **[45]** led to spindle collapse in 21% cells upon bipolar optogenetic activation. However, in the cells that successfully divided, spindle rotation was abolished (Fig. 3B), indicating that Dynein mediated pulling forces are essential. We next tested the importance of NuMA for cortical localization of Dynein by reducing its cortical levels using low doses of MLN-8237. This treatment results in partial inhibition of the activity of Aurora-A kinase, which leads to the re-localisation of NuMA from the cortex to the spindle pole **[46]** (Fig. S4A and B), blocking spindle rotation (Fig. 3B). This demonstrates an essential role for cortical NuMA for spindle rotation in response to bipolar GEF DH-PH activation. Finally, depolymerisation of astral microtubules using low doses of nocodazole that do not affect the spindle (Fig. S4C) also prevented spindle rotation (Fig. 3B), indicating that forces exerted by astral microtubules are essential for spindle rotation. Together, these experiments indicate that cortical regulators and astral microtubules participate in spindle rotation in response to bipolar GEF activation.

**Figure3:**
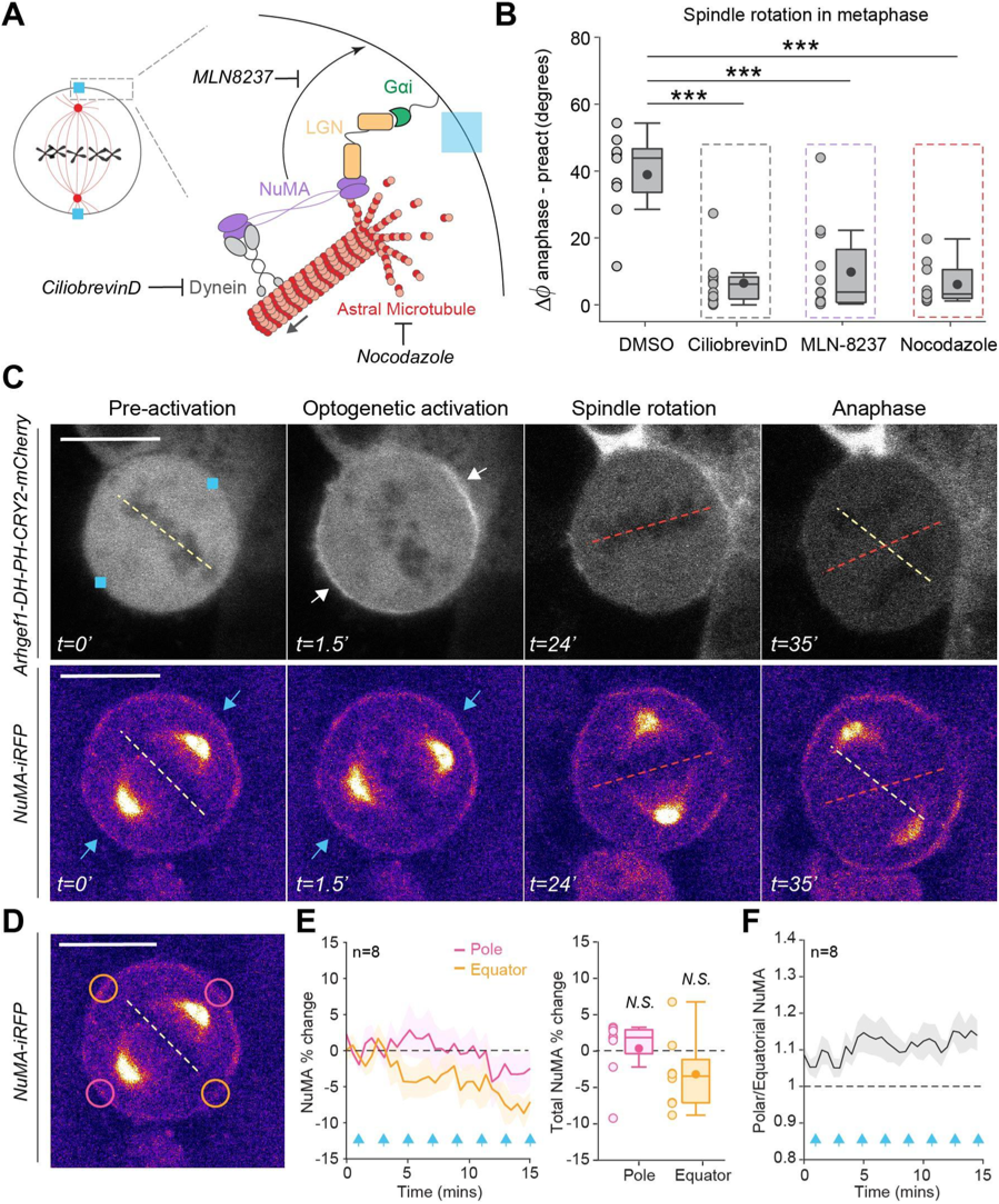
Spindle rotation depends on cortical localisation of NuMA and pulling forces exerted by Dynein. A. Schematic showing cortical regulators involved in spindle positioning. The ternary complex of Gαi, LGN and NuMA is anchored at the plasma membrane and recruits the motor protein Dynein. Dynein exerts minus end directed pulling forces on astral microtubules (grey arrow) to position the spindle. Ciliobrevin D blocks the ATPase activity of Dynein, low doses of MLN-8237 partially inhibit Aurora-A kinase, thus trapping NuMA at the spindle pole and blocking its transport to the cortex, and low doses of nocodazole block polymerisation of astral microtubules without severely affecting the spindle. B. Boxplot showing spindle rotation in metaphase for control cells treated with DMSO (n=9), 10μM ciliobrevinD (n=13), 100nM MLN8237 (n=12) and 20nM nocodazole (n=10). Boxplot shows median (dark line), interquartile range, mean (dark filled circle) and individual data points (light filled circle). ANOVA and Tukey-Kramer test: DMSO vs ciliobrevinD: *p* < 0.001 (***), DMSO vs nocodazole: *p*< 0.001 (***), DMSO vs MLN8237: *p*< 0.001 (***). C. Top: Time series showing localisation of Arhgef1-DH-PH-CRY2-mCherry in cells before and after bipolar optogenetic activation until anaphase onset. Bottom: Time series showing the corresponding localisation of NuMA-iRFP with low intensities appearing in black, medium intensities in magenta and high intensities at the spindle poles in yellow. Yellow and orange dashed lines indicate the position of the DNA in the metaphase plate before activation and at anaphase onset respectively. Blue squares (top) and blue arrows (bottom) indicate the region of activation, arrows point to regions of GEF-DH-PH accumulation (top panel) and indicate the same location in the NuMA channel (bottom panel). Scale bar = 10μm. Time is in minutes. D. NuMA-iRFP image at t=0 as in (C, bottom panel) showing the regions used to measure fluorescence intensity in (E-F), with pole (pink) and equator (yellow). Scale bar is 10μm. E. Left: Temporal plot showing the percentage change in NuMA-iRFP intensity at the pole and equator after bipolar activation. Plot shows mean ± SEM (n=8 cells). Blue arrows indicate the times of blue light stimulation. Right: Boxplot showing the total percentage change in NuMA-iRFP intensity at the pole and equator until 15 minutes post-activation for cells as in (E, Left). Boxplot shows median (dark line), interquartile range, mean (dark filled circle) and individual data points (light filled circle), student *t*-test compared to 0% change: Pole: *p*=0.8162 (N.S), Equator *p*=0.1162 (N.S). F. Temporal plot showing the ratio of polar/equatorial NuMA for cells as in (E).

We considered two possible mechanisms by which cortical regulators and astral microtubules could affect spindle rotation: first, changes in cortical tension following bipolar activation of GEF DH-PH may affect the localisation or enrichment of cortical regulators via mechanotransduction or second, spindle rotation may arise from a change in the balance of forces exerted on astral microtubules. Recent work has shown that optogenetic cortical targeting of NuMA in human cells is sufficient to recruit the dynein-dynactin complex and account for efficient spindle positioning **[47]**. Therefore, any change in the distribution of NuMA would lead to localised changes in the forces applied on astral microtubules and cause spindle rotation. To test this hypothesis, we imaged the localisation of iRFP-NuMA upon optogenetic activation. We observed no significant change in the localisation or intensity of NuMA fluorescence at the cortex over 14 minutes post bipolar activation (Fig. 3C, D and E), indicating that the localisation of cortical regulators contributing to pulling force generation is not perturbed. To verify that NuMA is polarised both before and after activation, we measured the ratio of polar/equatorial NuMA over time. This ratio remained above 1 throughout our experiment (Fig. 3F). We conclude that spindle reorientation in response to bipolar GEF activation does not occur due to a change in the localisation of cortical regulators. Instead, the spindle may reorient in response to changes in the cortex itself.

### Spindle rotation results from a local decrease in pulling force exerted on astral microtubules at regions of high cortical tension

To investigate our second hypothesis, we tested whether spindle rotation arises from a change in the balance of forces exerted on astral microtubules upon GEF DH-PH activation. First, we confirmed that metaphase spindles are subjected to a tensile force due to interaction of astral microtubules with the cortex **[48]**. To this end, we severed astral microtubules at one spindle pole using laser ablation and imaged changes in centrosome position. The rapid inward movement of the centrosome after ablation, indicated that the centrosomes are subject to a net pulling force (Fig. S5A and B).

Having established the resultant of the forces acting on the centrosomes in control conditions, we monitored the displacement of centrosomes in response to optogenetic activation. We reasoned that a displacement of the centrosome towards the cell centre would occur following a reduction in the cortical pulling forces and/or as a result of an increase in the microtubule pushing forces, while displacement away from the cell centre would suggest an increase in the pulling forces and/or a decrease in the pushing forces (Fig. 4A). First, we carried out optogenetic activation at only one pole (unipolar). This led to a displacement of the centrosome towards the cell centre on the activated side (activated pole), whereas the centrosome on the non-activated side (non-activated pole) underwent little if any change in position (Fig. 4B, C and D). Similarly, following bipolar activation, we observed an inward displacement of both centrosomes (Fig. 4C-top panel and E). These results suggest that optogenetic activation leads to either a reduction in the pulling force exerted by Dynein or to an increase in the pushing force exerted by astral microtubules in myosin-enriched tensed cortical regions.

**Figure4:**
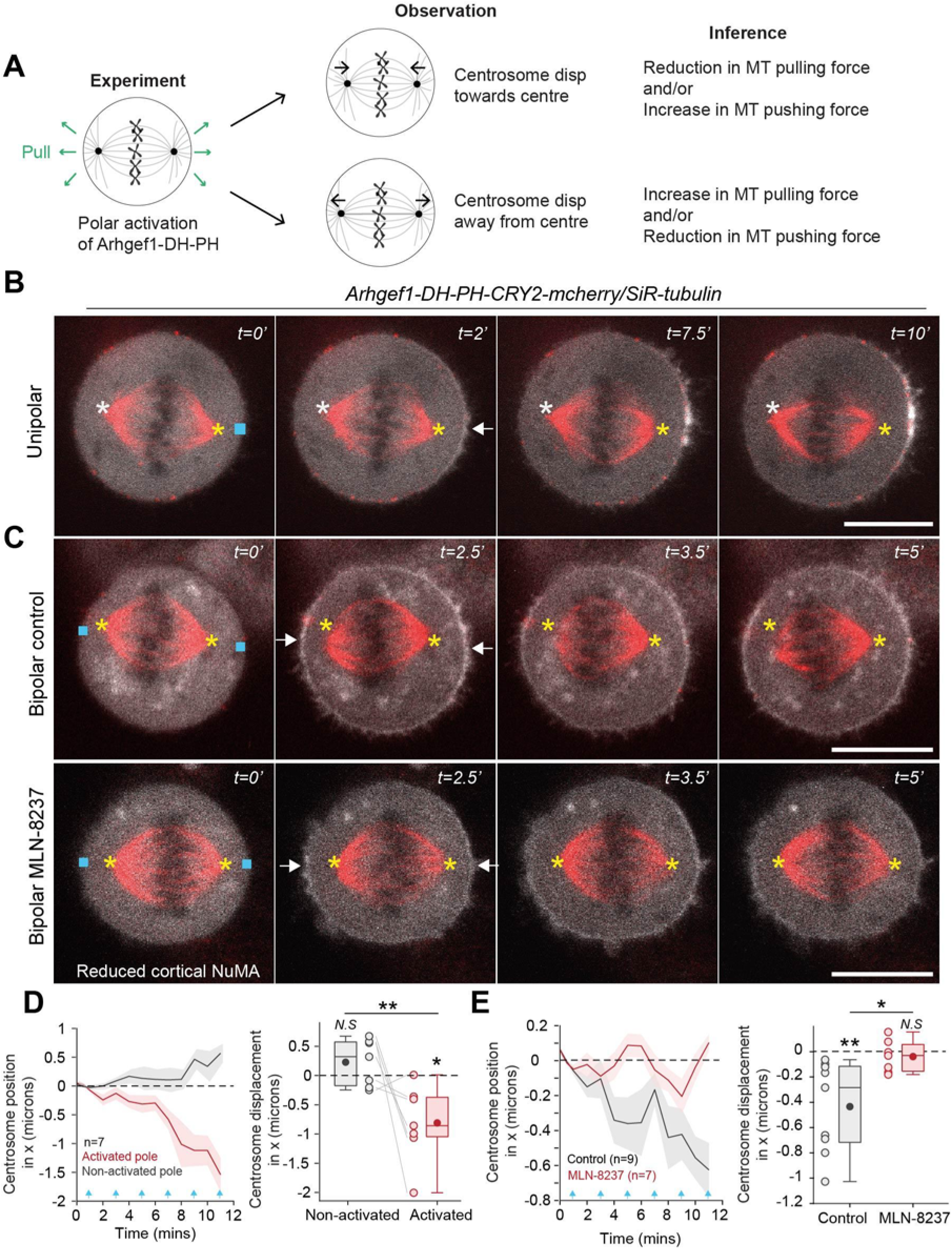
Spindle rotation results from a local decrease in pulling force exerted on astral microtubules at regions of high cortical tension. A. Schematic depicting the displacement of centrosomes as an indicator of forces acting on the spindle after polar optogenetic activation. Middle, top: An inward displacement of the centrosome towards the cell centre (black arrows) could result from a reduction in pulling force and/or an increase in pushing force. Middle, bottom: in contrast, an outward displacement of the centrosome away from cell centre (black arrows) would result from an increase in pulling force and/or a decrease in pushing force. B. Time series showing Arhgef1-DH-PH-CRY2-mCherry (greyscale) and siR-tubulin (red) in cells before and after unipolar optogenetic activation until anaphase onset. Blue square indicates the region of activation and the white arrow the region of GEF-DH-PH accumulation. The position of the centrosomes at t=0’ is marked as a yellow star (centrosome on the activated side-AP) and a white star (centrosome on the non-activated side-NAP). Scale bar = 10μm. Time is in minutes. C. Top: Time series showing Arhgef1-DH-PH-CRY2-mCherry (greyscale) and siR-tubulin (red) in cells before and after bipolar optogenetic activation until anaphase onset. Bottom: Time series as in the top panel in cells treated with MLN-8237 that have reduced cortical NuMA. Blue squares indicate the regions of activation and white arrows regions of GEF accumulation. The position of the two centrosomes at t=0’ is marked with yellow stars. Scale bar = 10μm. Time is in minutes. D. Left: Temporal plot showing the position of the two centrosomes along the x-axis following unipolar optogenetic activation for cells as in (B). The displacement of each centrosome is measured with the convention that movement towards the cell centre is negative and away from the centre is positive. Plot shows mean ± SEM (n=7cells). Blue arrows indicate the times of blue light stimulation. Right: Boxplot showing total centrosome displacement along the x-axis until 12 minutes (n=7cells). Student *t*-test compared to 0 μm displacement: NAP *p*=0.1813 (N.S), AP p=0.0169 (*); student’s *t*-test NAP vs AP *p*=0.0041 (**). E. Left: Temporal plot showing the position of the two centrosomes along the x-axis following bipolar optogenetic activation for control (n=9) and MLN-8237 (n=7) treated cells as in (C). The displacement of each centrosome is measured with the convention that movement towards the cell centre is negative and away from the centre is positive. Plot shows mean ± SEM. Blue arrows indicate the times of blue light stimulation. Right: Boxplot showing total centrosome displacement along the x-axis. Student *t*-test compared to 0 μm displacement: Control *p*=0.0069 (**), MLN-8237 p=0.4454 (N.S); unpaired student’s *t*-test Control vs MLN-8237: *p*=0.0156 (*).

To distinguish between these scenarios, we measured centrosome displacement in cells treated with MLN-8237, which have reduced levels of cortical NuMA and therefore less cortical Dynein (Fig. S4A and B) **[46]**. We reasoned that if optogenetic activation increases the pushing force, we should still observe centrosome displacement in MLN-8237 treated cells; whereas, if it reduces the pulling force, we should observe little or no centrosome displacement. Treatment with MLN-8237 abolished the inward displacement of the centrosomes (Fig. 4C-bottom panel, and E). Our data therefore suggests that localised changes in myosin abundance and cortical tension following bipolar GEF activation lead to a local reduction in the pulling force exerted on astral microtubules, resulting in centrosome displacement towards the cell centre. We hypothesize that it is this change in the balance of forces acting on the astral microtubules, that enables spindle rotation.

### Spindle rotation is an emergent property of the molecular mechanisms of force generation on astral microtubules

Under control conditions, the stable bipolar spindle orientation is established by forces exerted on astral microtubules by dynein motors, anchored to the plasma membrane by a complex comprising Gαi, LGN and NuMA. To explore how these forces result in a stable spindle position and how this stable position responds to changes in cortical tension, we developed a simplified mathematical model of spindle positioning based on our experimental data and current knowledge of the forces exerted on spindles (Supplementary theory). We consider a mitotic cell of radius *R* with two centrosomes separated by a distance *2l_c_*, from which astral microtubules emanate with a maximal length *l_m_* (Fig. 5A; S6A and B). The spindle’s orientation with respect to a horizontal reference axis is parametrised by an angle (*ϕ*) and the spindle is subjected to a torque (Γ) (Fig. 5A; S6A and B). In line with previous work, Γ arises from dynein motors exerting forces acting at the tip of astral microtubules touching the cell cortex **[49–52]**. The distribution of pulling forces along the cortex was taken to be proportional to the experimentally measured NuMA fluorescence profile (Supplementary theory; Fig. S4A and Fig. 5B). In the model, the cortical force distribution *f*(*θ*) was parametrised using a function of the form 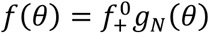, where 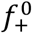 has a dimension of a pulling force per microtubule and *g_N_*(*θ*) is a periodic function whose shape is evaluated from the experimentally measured NuMA fluorescence profile along the cell periphery (Fig. S4A, Fig. S6E and Fig5B; Supplementary theory). With these data, our model computes a torque (Γ/Γ_o_) acting on the spindle as a function of its angular position (*ϕ*). Under control conditions (Fig. 5C, control), when *ϕ* is positive the torque is negative, signifying that the spindle will move back towards the 0° orientation and conversely, when *ϕ* is negative the torque is positive, again moving the spindle back towards 0°. Therefore, our model predicts that the 0° orientation is a stable position of the spindle under control conditions.

**Figure5:**
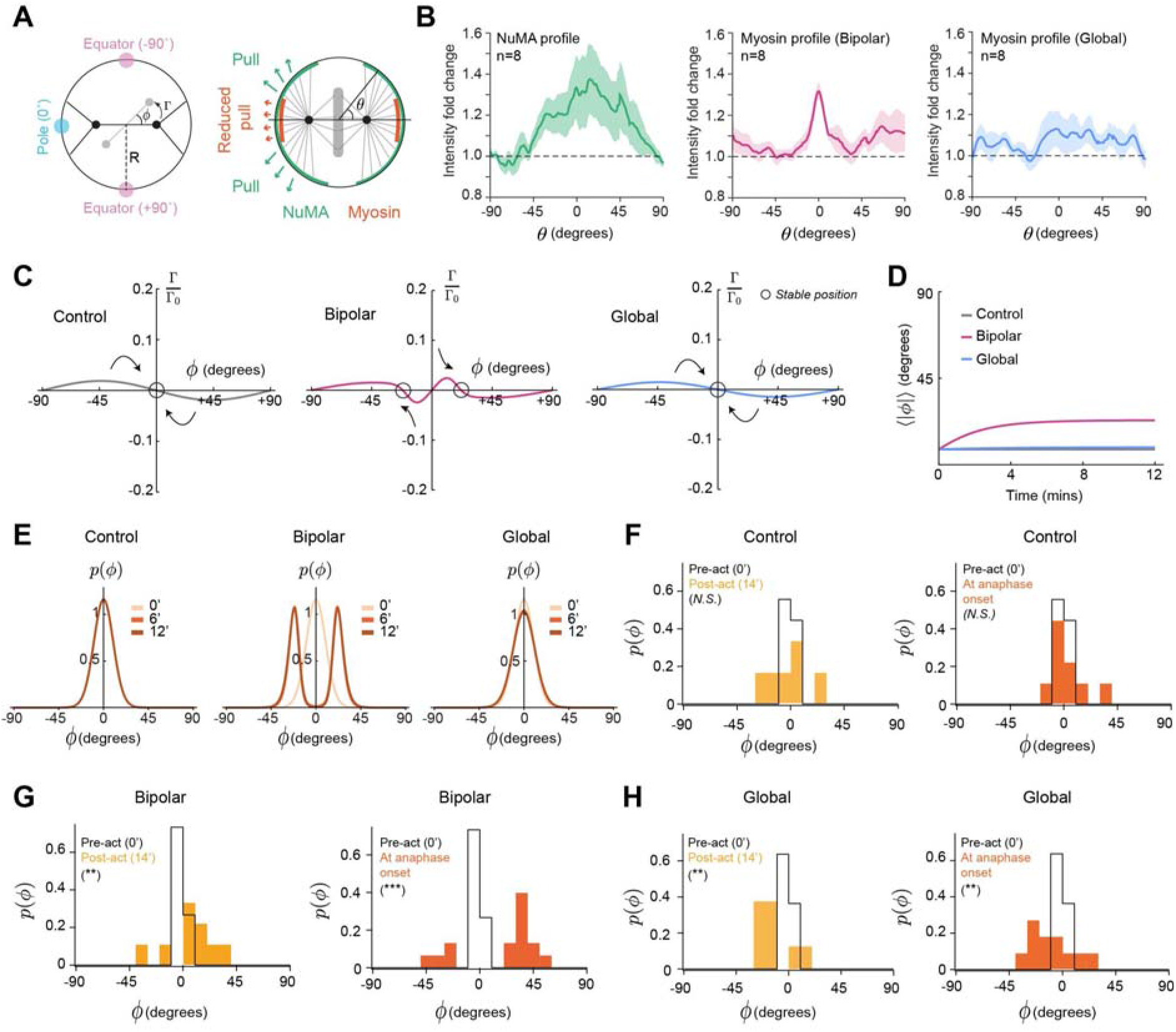
Spindle rotation is an emergent property of the molecular mechanisms of force generation on astral microtubules. A. Left: Schematic depicting the model of spindle positioning. We consider a circular cell of radius *R*. The spindle is depicted as a solid line linking two solid circles, which indicate the spindle poles. The reference spindle position is chosen to be horizontal and is shown in black. The spindle position at time t is shown in light grey. Astral microtubules emanate from the spindle poles with a uniform angular distribution and can contact the cortex if they are sufficient long (solid black lines). At time t, the spindle makes an angle *ϕ* with the horizontal axis, and is subjected to a torque Γ arising from cortical forces acting on astral microtubules. The spindle angle *ϕ* = 0 corresponds to the spindle pointing towards the cell poles, while *ϕ* = ±90° corresponds to the spindle pointing towards the equator. Right: Schematic showing the distribution of pulling forces at the cortex (green) as observed from the distribution of NuMA. *θ* denotes the polar angle of a point on the cell cortex. Astral microtubules are subjected to a pulling force when they interact with force generators at the cortex. In regions of optogenetic activation, they are subjected to a reduced pulling force (red). B. Left: Normalised fluorescence intensity cortical profile of NuMA along the cell cortex as a function of the angle *θ* obtained from experimental immunostaining data of NuMA. The profile is normalised to the intensity at the equator (n=8 cells). Centre: Spatial profile of fluorescence intensity of myosin along the cortex normalised to pre-activation intensity after bipolar optogenetic activation (n=8 cells), and Right: after global optogenetic activation (n=8 cells). *θ* = 0 represents the cell pole and *θ* = ±90° represents the equator. Plots show mean ± SEM. C. Predicted normalised torque (Γ/Γ_0_) as a function of spindle angle (*ϕ*) for control (Left), bipolar (Middle) and global (Right) activation. Arrows indicate the direction of spindle rotation. Stable spindle positions are indicated by black circles. Profiles of cortical pulling forces and reduced cortical pulling forces are taken proportional to the fluorescence intensity profile of NuMA and myosin (B). See supplementary theory for further information. D. Predicted average value of the spindle orientation as a function of time after optogenetic activation for different conditions. The spindle orientation is measured between −90 ° and 90°, and the absolute value is averaged. E. Predicted probability distribution of the spindle angle, *p*(*ϕ*), at successive times (t=0,6,12 mins) after optogenetic activation for control, bipolar and global activation. F. Histogram showing the experimental probability distribution of the spindle angles, *p*(*ϕ*), plotted as a function of spindle angle (*ϕ*), pre-activation and 14 minutes post-activation (Left) and pre-activation and at anaphase onset (Right) for control cells. 2-sample Kolmogorov-Smirnov test, Control pre-act vs 14’: *p*=0.23 (N.S), pre-act vs anaphase onset: *p*=0.60 (N.S). n=9 for pre-act and anaphase onset; n=6 for 14’ post-activation. G. Histogram, as in F, for cells after bipolar activation. 2-sample Kolmogorov-Smirnov test, Bipolar pre-act vs 14’ *p*=0.0065 (**), pre-act vs anaphase onset *p*=2.3×10^-4^ (***). n=15 for pre-act and anaphase onset; n=9 for 14’ post-activation. H. Histogram, as in F, for cells after global activation. 2-sample Kolmogorov-Smirnov test, Global pre-act vs 14’ *p*=0.0046 (**), pre-act vs anaphase onset *p*=0.0025 (**). n=11 for pre-act and anaphase onset; n=8 for 14’ post-activation.

We next asked how the spindle position would change under a theoretical perturbation of the distribution of cortical forces. We found that, when the spindle is subjected to an additional bipolar profile of cortical forces locally reducing the pulling force, the spindle can either keep its stable 0° orientation or leave its original position to move to a new stable orientation, depending on the magnitude of the perturbation and whether net pushing forces can arise (Fig. S6F, G and H). Next, we parameterised the cortical force distribution for our optogenetic activation experiments. Since the distribution of NuMA does not change with optogenetic activation (Fig. 3C, D, E and F), *g_N_*(*θ*) was kept the same as in the control condition. Additionally, the change in myosin distributions for both bipolar and global activation conditions were fitted with the periodic function *g_M_*(*θ*) (Fig. 5B, S6C and E; Supplementary theory). The force distribution acting on the spindle was then computed by adding a new contribution to the force profile, now taking the form 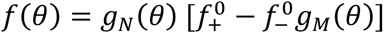, where *g_M_*(*θ*) is proportional to the change in myosin intensity after activation. Here, based on our experiments, we assume that the effect of myosin is to decrease cortical pulling forces on microtubules, with a magnitude that depends both on the NuMA and the myosin distributions. To simulate decreased pulling forces following activation, we chose values of the ratio, 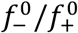, that were sufficiently large but still low enough such that *f*(*θ*)>0 (Fig. S6I). Our model predicted a new stable spindle orientation for bipolar activation but no change for global activation, qualitatively consistent with our experiments (Fig. 5C, bipolar and global). Indeed, under bipolar activation conditions, when *ϕ* is positive, the torque is positive, indicating that the spindle will move away from 0°, towards a new, positive tilted orientation. Conversely, when *ϕ* is negative the torque is negative indicating that the spindle moves away from 0° and towards a negative tilted orientation. Therefore, our model predicts that mitotic spindles move away from the 0° orientation to a new stable position in response to bipolar activation but not in response to global activation.

Next, we asked if our simulation could capture the dynamics of rotation by using a dynamical model of spindle motion. Here, the spindle moves according to the torque acting on it and its movement is resisted by an effective rotational friction force. Our experimental data indicated that, even under control conditions, the spindle angle tends to fluctuate (Fig. S6D). We modelled these fluctuations by introducing a random force acting on the spindle motion, leading to spindle diffusion with a diffusion constant *D*. Experimental measurements of the mean square rotational displacement of the spindle then allowed us to determine the diffusion constant *D* and a characteristic time 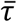, that depend on friction and on the pattern of cortical forces (Supplementary theory and Fig. S6D). This, in turn, allowed us to predict the temporal evolution of the spindle orientations. The predicted time that the spindles needed to reach their stable orientation was roughly consistent, although slightly shorter when compared to the median time until anaphase onset (~15 minutes) from experiments (Fig. 5D, Fig.2I, Fig. S6J and K).

Increasing the magnitude of the ratio 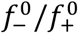 led to spindle dynamics closer to those observed in experiments, where the spindle position did not saturate by 15 minutes (Fig 1L and Fig S7C). In this case, the model predicted that both bipolar and global optogenetic activation would give rise to a new stable spindle orientation (Fig. S7A). The kinetics of reorientation were slower with spindles reaching their stable orientation only after ~40 minutes, much longer than the median time until anaphase onset in our experiments (~15 minutes, Fig. S7E and F and Fig. 2I). When we limited our predictions to durations of 15 minutes, bipolar activation gave rise to rotations consistent with those observed experimentally (Fig. S7C and D), whereas global activations only gave rise to small rotations because of the lower torques resulting under this condition (Fig. S7A, C and D). However, from a mechanistic point of view, this value of 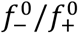 implied that interaction of microtubules with the cortex resulted in net pushing forces (Fig. S7B), in contrast to the situation examined previously in which only pulling forces act on the spindle (Fig. S6I). Given that we do not observe inward movement of the centrosomes in response to optogenetic activation in the presence of MLN-8237 (Fig. 3C and E), and that spindle rotation is abolished following treatment with ciliobrevin D (Fig. 3B), such pushing forces would have to be dependent on the activity of NuMA and dynein, a mechanism not supported by current knowledge of spindle positioning.

The model allowed us to predict not only the average position of the spindle, but also the probability density function *p*(*t, Φ*) of spindle orientation at time *t*. To test how this probability changed with a given perturbation, we plotted the probability density function from the model for our three conditions at different times (Fig. 5E) and compared these to the experimentally observed probability distributions (Fig. 5F, G and H). We found that, under control conditions, the distribution of spindle orientations remains steady over time and is peaked around 0° (Fig. 5E, control). During the 15 minutes that correspond to the median time until anaphase onset in experiments, *p*(*t,Φ*) splits into two symmetric peaks over time in response to bipolar activation, whereas *p*(*t,Φ*) retains a single peak that broadens slightly over time in response to global activation (Fig. 5E, bipolar, global). These model predictions were in good qualitative agreement with our experimental probability distributions, computed pre-activation, 14 minutes post-activation and at anaphase onset for all three conditions (Fig. 5F, G and H). However, the distribution of spindle orientation in response to global activation displayed more extensive broadening under our experimental conditions than in simulations.

Overall, our model predicts that a reduction of pulling force in myosin-enriched regions can cause the spindle to become unstable, rotating away from its initial stable position. Taken together, our model and experiments argue that spindle orientation can be understood from the interplay between cortical mechanics and the spindle positioning machinery.

## Discussion

In this study, we have shown that mitotic spindles can change their orientation in response to inhomogeneities in myosin abundance and cortical tension, thereby affecting the axis of cell division. In our experiments, we increase cortical tension by activating RhoA via optogenetic re-localisation of GEF DH-PH. Recruitment of the GEF specifically to polar cortical regions in metaphase leads to myosin enrichment in these regions, a local increase in cortical tension, and subsequent rotation of the spindle away from cell poles. Whereas pulling forces mediated by cortical NuMA/Dynein are essential to power rotation, their distribution does not change upon GEF recruitment. Instead, our data suggests that a local reduction in the pulling force exerted on astral microtubules in tensed, myosin-rich regions destabilises the spindle from its original position. The larger pulling force exerted by NuMA/Dynein outside the region of activation generates a torque on the spindle leading to its rotation. A mathematical model incorporating this change in the force balance can qualitatively predict the spindle orientation distributions and rotation dynamics observed in experiments. Taken together, our data suggest that spindle reorientation in response to localised mechanical changes in the cortex is an emergent property of the interaction between the cortex and the spindle positioning machinery (Fig. 6).

**Figure6:**
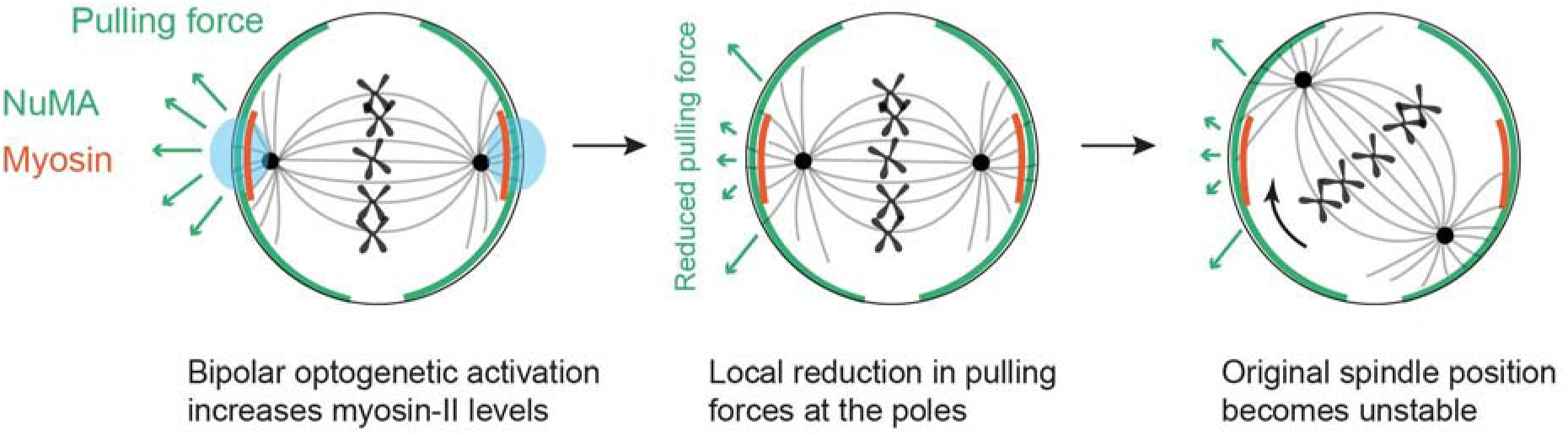
A schematic showing spindle rotation after bipolar Arhgef1 DH-PH activation. Schematic depicting the biophysical mechanism of spindle rotation observed after bipolar recruitment of Arhgef1-DH-PH in metaphase. Left: The spindle is initially in a stable position controlled by pulling forces exerted on astral microtubules (green). Optogenetic activation at the poles (blue) results in local enrichment of myosin at these regions (red). Middle: Increased tension at the poles generated by myosin activity leads to a local reduction in the pulling force exerted on astral microtubules. Right: The original spindle position becomes unstable. Indeed, the distribution of NuMA at the cortex is broader than the activated region and not perturbed after activation, therefore pulling forces powered by Dynein/NuMA will exert a torque on the spindle causing spindle rotation. The spindle rotates towards its new stable position and the cell undergoes anaphase and cytokinesis.

### Spindle reorientation is an emergent property of the interplay between cortical mechanics and the spindle positioning machinery

Our model and experiments suggest that spindle rotation in response to a change in RhoA activity and cortical tension represents an emergent property of the interaction between the spindle positioning machinery and the cortex, rather than resulting from mechanotransductory signalling pathways. Indeed, our optogenetic activation experiments along with our laser ablation data indicate that a localised increase in cortical tension caused by myosin recruitment at the poles decreases pulling forces on the centrosomes. Our theoretical model can predict spindle rotation away from its initial stable position, simply by assuming that the experimentally measured myosin activation profile results in a reduction of cortical pulling forces. Although our current model is in qualitative agreement with our experiments, the magnitude of the rotation it predicts is smaller than our observations. One likely source for this discrepancy is that, although our experiments indicate a reduction of pulling forces in myosin-rich tensed regions, we currently lack a detailed characterisation of how myosin contractility and tension impinge on the profile of pulling forces generated by dynein. In line with our experiments, our simulations only incorporated net reduction in pulling forces; however, if net pushing forces are present, our model indicates that spindles would rotate to a new stable position of 90° rather than ~20° (Supplementary theory and Fig. S7A). Interestingly, the torque and velocity of rotation depended on the relative magnitude of pushing forces compared to pulling forces. When pushing forces were comparable or larger than pulling forces, rotation was slower and the time necessary to reach steady-state orientation became several-fold larger than the median time until anaphase onset, potentially limiting how far spindles can rotate away from their initial orientation (Supplementary theory and Fig. S7B, C, D and E). It will be interesting to investigate the applicability of our model to organisms in which microtubule pushing has been evidenced, such as *C.elegans* **[41]**.

How tensed, myosin-enriched cortical regions generate less pulling force on astral microtubules is not understood. In addition, which of myosin enrichment or increased tension causes reorientation remains unclear. Although blebbistatin treatment prevents myosin force generation, it does not prevent myosin binding strongly to F-actin **[53, 54]**. As blebbistatin abolished spindle rotation, this suggests that spindles do not respond to myosin enrichment alone but likely to a change induced by myosin force generation and increased cortical tension. An increase in myosin contractility and cortical tension may lead to a more rigid cortex causing a reduced efficiency of force generation by dynein, and/or decrease the lifetime of dynein-astral microtubule interaction, thereby reducing the average pulling force per astral microtubule. Alternatively, myosin contractility and increased cortical tension at the poles may change cortex architecture, for example by reducing the average size of gaps within the cortical F-actin mesh. Such a reduction in mesh size may prevent astral microtubules from reaching the plasma membrane where the Gαi-LGN-NuMA-Dynein complex is located, again leading to a reduction in the average pulling force per astral microtubule. Future experiments will be necessary to determine the biophysical mechanism through which cortical tension and/or myosin enrichment reduce pulling forces.

### Inhomogeneities in cortical tension control spindle orientation

Previous work has revealed that spindle orientation is sensitive to dynamic changes in the stress applied to dividing cells **[14, 17, 20].** However, little is known about the subcellular mechanical changes that cause reorientation. In our experiments, spindle reorientation was induced by a localised increase in RhoA activity at the poles, which led to a localised myosin contractility and a localised increase in cortical tension. Polar increase in myosin contractility leads to a localised increase in cortical tension giving rise to a change in the cellular aspect ratio, a cue known to influence the orientation of cell division **[55]**. However, bipolar activation led to only a small deformation of ~3% (Fig. S1B and C). Moreover, in our experiments, we focused on spherical mitotic cells (aspect ratio <1.2) to examine the impact of inhomogeneous cortical tension on spindle orientation in isolation from shape cues. Therefore, we suggest that spindles likely sense inhomogeneities in cortical tension possibly directly because of differences in myosin activity, and rotate away from regions of high tension with small contribution from cell shape. In future, it will be interesting to dissect the competition between shape and inhomogeneous cortical tension in the control of spindle orientation.

In living tissues, tension can arise from either active or passive processes. Active stress originates from the action of myosin motors on the cell cytoskeleton, whereas passive stress arises from deformation of cytoskeletal networks in response to external forces. While in our experiments, cortical tension inhomogeneities were generated through differences in active stress due to myosin enrichment, spindles have also been shown to reorient in response to passive stress arising from application of a uniaxial stretch to isolated cells and tissues **[14, 17]**. Isolated metaphase cells seeded on elliptical micropatterns were stretched along their short axis, rendering the cells spherical. Spindles that were initially oriented along the long axis of the cell reoriented with an angle of ~40° in ~15-20 min before entering anaphase, similar to our experiments **[17]**. Similarly, in the zebrafish enveloping layer, application of an ectopic tissue stress perpendicular to the long axis of a dividing cell resulted in spindle reorientation with an angle of ~30° within 6-7 minutes **[14]**. In both studies, the likely result of mechanical manipulations is to increase cortical tension at the poles of the mitotic cell. Interestingly, the dynamics and magnitude of spindle reorientation in these studies is comparable to our data, suggesting that reorientation may be independent of the exact process through which cortical tension inhomogeneity is generated.

### Physiological consequence of spindle reorientation

Reorientation of mitotic spindles in response to a sudden application of stress may help optimise cell packing in tissues subjected to deformations as part of their physiological function **[56]** or in organ development where mechanical stresses play an integral part in guiding further morphogenesis **[57–59]**. While previous studies have examined the orientation of cell divisions in response to a constant stretch lasting several hours **[6, 14, 23]**, many epithelial tissues such as the skin, bladder, or the intestine are subjected to transient deformations, which generate transient inhomogeneous stress fields. Our study indicates that, despite their transient nature, these stresses may be sufficient to reorient the axis of division of metaphase cells. Such a phenomenon may allow the tissue to optimise its organisation to reduce stress in the direction of extension.

## Data availability

Data and analysis codes will be made available online upon publication on the UCL data repository (https://rdr.ucl.ac.uk/) with a unique doi (10.5522/04/16871626).

## Acknowledgments

We thank Susana Godinho (Queen Mary University of London, UK) for gateway cloning plasmids, Mathieu Coppey (Institut Curie, Paris, France) for advice on optogenetics and sharing reagents, Ayad Eddaoudi (UCL, Institute for Child Health) for flow cytometry analysis, Jonathan Fouchard, Julia Duque, Kazunori Yamamoto and Diana Khoromskaia for comments and discussions, and members of the Charras and Baum labs for discussions. MK was supported by a SNSF early post-doc fellowship P2LAP3_164919 and by a European Research Council consolidator grant (CoG-647186) to GC. PB was supported by a CRUK multidisciplinary award (C55977/A23342) to GC and GS. GS was supported by the Francis Crick Institute which receives its core funding from Cancer Research UK (FC001317), the UK Medical Research Council (FC001317) and the Wellcome Trust (FC001317).

## Author contributions

M.K., G.S. and G.C. conceived the project, wrote the manuscript and secured the funding. M.K. performed most of the experiments and analysis. P.B. performed the AFM experiments and analysis. M.S. provided the plugin for segmentation analysis. V.S. contributed to analysis and discussions. A.L. advised on protocols, analysis and discussions. E.F. and L.V. helped in flow cytometry and generation of plasmids. B.B. advised on experiments and discussions. G.S. implemented the model and advised on analysis. G.C. helped oversee the entire project and writing. All authors discussed the results and manuscript.

## Supplementary information

### Materials and Methods

#### Cell Lines and Culture

MDCKII cells were cultured at 37°C with 5%CO_2_ in high-glucose (4.5g/L) DMEM (Thermo Fisher) supplemented with 10% fetal bovine serum (Sigma), 1% penicillin-streptomycin (Thermo Fisher) and 25mM Hepes buffer (Gibco). Where appropriate, the medium was supplemented with selection antibiotics: G418 1mg/mL (Sigma), puromycin 1μg/mL (Calbiochem) and HygromycinB 300μg/mL (Invivogen). A stable cell line expressing CIBN-GFP-CAAX was made by retroviral transduction into MDCKII WT cells. Cell lines stably expressing CRY2-mCherry, ARHGEF1-CRY2-mCherry, MRLC-iRFP, LifeACT-iRFP or iRFP-RhoA biosensor were made by lentiviral transductions. A stable cell line expressing iRFP-NuMA was made by electroporation (Lonza). All cell lines were selected with appropriate antibiotics and sorted by flow cytometry before use. Cells were routinely tested for the presence of mycoplasma using the mycoALERT kit (Lonza).

#### Cloning

The plasmids pCRY2PHR-mCherryN1 and pCIBN(deltaNLS)-pmGFP were acquired from Addgene (plasmid #26866, and plasmid #26867), **[1]** The DH-PH domain of ARHGEF1 (p115-RhoGEF) was identified using Uniprot. We extended the sequence of interest to retain 8 extra amino acids at either extremity of this catalytic domain following a previously used approach **[2].** This domain was synthesized by GenScript (New Jersery, United States) and inserted into the CRY2-mCherry plasmid using NheI/XhoI cloning sites. The obtained plasmid ARHGEF1-DH-PH-CRY2PHR-mCherryN1 was then inserted into a lentiviral vector using the Gateway technology first into a *p-DONR-221* vector and finally in a *p-DEST*-Neomycin vector (Thermo Fisher). CIBN(deltaNLS)-pmGFP was inserted in a retroviral vector pRetroQAcGFP-N1 (Clontech). MRLC-iRFP and LifeACT-iRFP plasmids were a kind gift from Leo Valon (Institut Pasteur, Paris, France). iRFP-Anillin-AHD-PH (RhoA biosensor) plasmid was made by cloning the Anillin-AHD-PH domain from the Addgene plasmid (#68026) into an iRFP-C1 plasmid. iRFP-C1 was made by replacing the eGFP with iRFP in the Clontech plasmid eGFP-C1 using AgeI/BsrGI restriction sites. MRLC-iRFP, LifeAct-iRFP, and iRFP-Anillin-AHD-PH were also inserted into a *p-DEST*-hygromycin lentiviral vector using Gateway technology. All clones were verified by sequencing (SourceBioscience). All plasmids contain the generic CMV promoter from the backbone pmCherry-N1 (Clontech Laboratories).

#### Sample preparation for microscopy

For all optogenetic experiments (except Fig1B, C), ~50,000 cells were plated on 35mm glass bottom dishes (World Precision Instruments) ~36hours prior to imaging. Cells were synchronised in pro-metaphase by treatment with nocodazole (30ng/mL) for 2 hours after which they were washed thrice with PBS to remove the nocodazole. Cells were then incubated with fresh phenol free medium for 10-15 mins at 37°C and 5%CO_2_ to release them in metaphase. All imaging experiments were carried out in phenol red free DMEM (ThermoFisher) supplemented with 10% fetal bovine serum (Sigma). In experiments to visualize microtubules (Fig4B, C and S5A), 50nM siR-tubulin dye (Spirochrome) was added 2 hours prior to imaging.

#### Confocal imaging and Optogenetic activation

Cells were imaged at 37°C in a humidified atmosphere with 5%CO_2_. Rounded metaphase cells were first identified using a 60x oil immersion objective (NA 1.35, Olympus) and all activation and imaging experiments (except in Fig3C and FigS3A) were carried out using a 100x oil immersion objective (NA 1.40, Olympus) mounted on an Olympus IX83 inverted microscope equipped with a scanning laser confocal head (Olympus FV-1200). Imaging in Fig3C and FigS3A was carried out using a 60x oil immersion objective (NA 1.35, Olympus). For brightfield imaging, a far-red filter was inserted into the illumination path to avoid spurious activation of CRY2. All optogenetic activation experiments (except in Fig1B, C) were carried out on Olympus FV-1200, using the 473-nm light set at 2% laser power (~0.8mW). Defined regions of interest were illuminated for 348 msec every 2 mins until the end of cell division. For polar and equatorial activation, a square region of 0.62μm x 0.62μm was illuminated whereas for global activation the whole field of view was illuminated using 473-nm light. Images were acquired in the medial focal plane of the dividing cell with a 30sec time interval until the end of cell division. Imaging of CRY2-mCherry was performed using a 559-nm laser and imaging of MRLC-iRFP, LifeACT-iRFP, iRFP-RhoA biosensor and iRFP-NuMA was done using a 633-nm laser. A GaAsP-high sensitivity detector unit was used to image mCherry and iRFP signals and reduce laser exposure times.

#### Sample preparation for cortical tension measurements

Cortical tension measurements (Figure1B,C) were carried out on round interphase cells as described in **[3].** Briefly, one hour before the experiment, cells were detached using trypsin-EDTA, resuspended in Leibovitz’s L-15 medium (Life Technologies) supplemented with 10% fetal bovine serum (Sigma) and plated onto 35-mm glass bottom dishes (World Precision Instruments) at low density to obtain single rounded cells.

#### Cortical tension measurements

Tension measurements were performed using a tipless silicon cantilever (ARROW-TL1Au-50-NanoWorld) with a nominal spring constant of 0.03N/m mounted on a JPK CellHesion module (JPK instruments) on an IX81 inverted confocal microscope Olympus-FV1000 (Olympus). Imaging was done with a 63x oil immersion objective (Olympus, N.A.=1.42). Sensitivity was calibrated by acquiring a force curve on a glass coverslip and the spring constant was calibrated by the thermal noise fluctuation method. The spring constant estimated for each experiment ranged between 0.055 and 0.081N/m. A first force-distance curve was acquired on a glass surface close to the cell to determine the distance between the retracted cantilever and the glass surface (hcantilever). A second, force-distance curve was acquired on top of the cell allowing determination of the cell height (h) relative to h_cantilever_. As the cantilever is attached to a glass block with a 10° tilt, the cell height prior to compression was obtained by measuring the distance (d) between the centre of the cell and the tip of the cantilever: h _cell_ = h+ d tan10°

The approach speed of the cantilever was set at 10μm/s. We chose a set-point force of 20nN, which produced an average cell compression of 2.5 - 4.7μm, which is 24-35% of the cell height. During the constant height compression, the force acting on the cantilever was recorded. After initial force relaxation of ~40sec, the resulting force value was used to extract surface tension (pre-activation) (γ_pre activation_). At about ~80sec after the onset of compression, optogenetic activation was carried out on Olympus FV-1000, using the 488-nm light at 10% laser power (2.3mW)._A circular region encompassing the whole cell was illuminated for 1 sec and the resulting force at ~120sec_after activation was used to determine the surface tension (post-activation) (γ_post activation_). The calculation of cortex tension is based on **[4, 5]** Briefly, neglecting difference in height of compression across the top surface of the cell due to the angle between the cantilever and the horizontal, **[3],** and assuming negligible adhesion between cell, dish and cantilever, the surface tension γ can be calculated as:

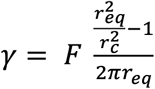

where r_eq_ is the equatorial radius of the selected cell, r_c_ is the radius of the contact between the cell and the cantilever, and F is the force exerted by the cell on the cantilever. r_eq_ was extracted using the level set method by detecting the contour of the cell in confocal images acquired after optogenetic activation **[6].** As we cannot have a direct measurement of r_c_, the contact radius was calculated using the following formula **[5]:**

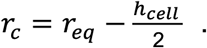

#### Laser ablation

Laser ablation of astral microtubules in FigS5A, was carried out on WT MDCK cells stained with SiR-tubulin dye to visualise microtubules. As it is difficult to visualise astral microtubules in live cells using these dyes, we chose a small ablation region between the spindle pole and the cortex, reasoning that astral microtubules should be present in this region based on our immunostaining images (FigS4C). A circular region of (0.496 μm x 0.496 μm) was exposed to a pulsed 405-nm laser (PicoQuant) at 100% power for 20sec delivered through a FV-1200 scanning laser microscope (Olympus) and imaging was carried out using a 100x oil immersion objective (NA, 1.40 Olympus) for 20mins post ablation.

#### Drug treatments

Drug treatments were performed as follows. To inhibit myosin contractility, s-nitro-blebbistatin (Cayman Chemicals) was added at 20μM concentration 10mins before experiments. We used this more stable form of blebbistatin as it has been reported that addition of a nitro group stabilizes the molecule thus preventing its degradation when exposed to 473-nm light **[7].** To block Rho-kinase activity, Y-27632 (Merck) was used at 50μM concentration 10mins before experiments. To inhibit Dynein, ciliobrevinD (Calbiochem) was added at 10μM 10mins before experiments **[8].** At this concentration, we could observe collapse of spindles in some cells and these were excluded from analysis. To inhibit polymerisation of astral microtubules without significantly affecting the spindle, we treated cells with low doses of nocodazole (20nM) for 10mins before experiments. To prevent cortical localisation of NuMA, we partially inhibited AuroraA activity using low doses of MLN-8237 (100nM) for 10mins before experiments **[9].**

#### Immunostainings

For immunostaining of NuMA in FigS4A, cells were pre-extracted in PHEM buffer (PIPES-18.14g, HEPES-6.5g, EGTA-3.8g, MgSO_4_-0.99g dissolved in 500ml H_2_0 adjusted to pH 7.0 with KOH) containing 0.3% Triton X-100 for 4min, fixed with 4% formaldehyde in PHEM buffer for 10-15min, permeabilised with 0.3% Triton X-100 in PHEM buffer for 5-10min, blocked in 3% BSA + 0.05% Tween-20 in PBS for 1 hour, all at room temperature (RT). Cells were then incubated overnight at 4°C with a rabbit monoclonal primary antibody against NuMA (Abcam-ab109262, 1:200 dilution) followed by three washes with PBS+3%BSA+ 0.05% Tween-20 each lasting 5 min. Cells were later incubated with a goat anti-rabbit secondary antibody conjugated with Alexa 488 (ThermoFisher, 1:200 dilution) for 4-6 hours at RT. This was followed by three washes with PBS+3%BSA+ 0.05% Tween-20 each lasting 5 min and staining of nucleic acids with Hoechst 33342 (5μg/mL-Merck Bio-sciences) for 5min. Following staining, cells were mounted in FluorSave reagent (Calbiochem) and imaged on Olympus FV-1200 using a 100x objective (NA 1.40, Olympus).

For immunostaining of microtubules in FigS4C, cells were fixed in ice-cold methanol at −20°C for 15min, followed by three washes in PBS each lasting 5min, blocked in PBS+10% horse serum (HS) for 5min 3x and incubated with a mouse monoclonal primary antibody against α-tubulin (Abcam DMA1, 1:1000 dilution) for 1 hour at RT. This was followed by three washes in PBS+10%HS for each lasting 5min, incubation in goat anti-mouse secondary antibody conjugated with Alexa 488 (Thermofisher, 1:200 dilution) for 1hour at RT, three washes in PBS+10%HS for each lasting 5min and staining of nucleic acids with Hoechst 33342 (5μg/mL-Merck Bio-sciences) for 5min. Following staining, cells were mounted in FluorSave reagent (Calbiochem) and imaged on Olympus FV-1200 using a 100x objective (NA 1.40, Olympus).

#### Determination of cell shape

For determination of cell shape in FigS1B,C, a line was drawn manually in Fiji along the spindle pole axis (polar axis) and along the DNA axis (equatorial axis) pre-activation and 2 minutes post-activation. The ratio of polar/equatorial axis was measured and the difference in this ratio (post-activation minus pre-activation) was measured as the Δ ratio.

#### Determination of spindle rotation angle and time until anaphase onset

To measure spindle rotation angle in Fig1J, FigS2G,H,J a line was drawn manually in Fiji i) through the metaphase DNA plate on the first frame before optogenetic activation, ii) in between the segregating chromosomes on the first frame at anaphase onset and iii) in between the 2 daughter cells at cytokinesis. The angle measured in all cases was the angle between the line and the horizontal. Spindle rotation in metaphase was defined as the absolute difference between the angle at anaphase onset and pre-activation; and spindle rotation in anaphase was defined as the absolute difference between the angle at cytokinesis and anaphase onset. The time until anaphase onset was taken as the difference in time between the first frame at anaphase onset and pre-activation.

#### Determination of spindle rotation speed

To measure the spindle rotation speed in Fig1K, the angle reached at anaphase onset was divided by the total time each cell took to reach anaphase onset.

#### Determination of spindle position over time

To measure the spindle position over time in Fig1L and S2K, the absolute spindle angle was measured by drawing a line manually in Fiji passing through the metaphase DNA plate every 1 minute before and after activation. The average spindle position was taken as the mean of the spindle angles between 11-14 minutes.

#### Determination of centrosome position and displacement

Centrosome position in Fig4D,E and S5B was measured as follows. All images were rotated such that the spindle was parallel to a horizontal line. The position of the centrosome stained with SiR-tubulin dye was manually tracked in Fiji and the x and y coordinates were obtained every minute until anaphase onset. A mean position for the pre-activation frames was calculated and the position in each frame was subtracted by this mean value for normalisation. Centrosome position along the x-axis was plotted as mean± SEM by converting the x-coordinate value from pixels to microns. To plot centrosome displacement, an average of the centrosome position for the frames pre-activation (X_pre_) and an average of the last 5 minutes post-activation (X_post_) was calculated. Centrosome displacement was obtained as X_post_ - X_pre_ for individual cells.

#### Bleach correction and Image registration

All images were corrected for photobleaching with the in-built Fiji plugin, using exponential curve fitting. Image registration was carried out using a custom-written spatial cross-correlation method implemented on MATLAB. The pre-activation frames of the bleach-corrected CRY2 image sequence were averaged to obtain a reference image. Each frame of the CRY2 image sequence was then spatially cross-correlated to the reference image to determine the spatial position where the cross-correlation was maximum. The frame was then offset along the X and Y axes according to the offset values shown by the cross-correlation peak. The offsets obtained, on a given frame, for the CRY2 channel were also applied to the other channel.

#### Temporal analysis of CRY2, LifeACT, Myosin, RhoA and NuMA fluorescence intensity

To measure the percentage change in fluorescence intensity of CRY2-GEF-mCherry, CRY2-mCherry, MRLC-iRFP, LifeACT-iRFP, iRFP-RhoA biosensor and iRFP-NuMA in Fig1 E, G, I, S2B-F, 2C-D, S3A,B and 3E the following method was used. Images were corrected for photobleaching and registered to correct for XY drift as described above. Circular ROIs (30×30 pixels) corresponding to the pole, equator and future pole wherever appropriate, were manually drawn in Fiji and the mean intensity in this region was obtained over time to get a temporal intensity profile. Fluorescence intensity was corrected for background by drawing the same circular ROI outside the cell. An average of the fluorescence intensity at the pole, equator and future pole in the frames pre-activation (F_0_) and the frames post-activation (F) was obtained. The percentage change in fluorescence intensity was plotted as (F-F0)/F0 x100 for each cell and the mean ±SEM was shown in the temporal plot. To measure the total change in fluorescence intensity, the percentage change in intensity of 22 frames (11 minutes) post-activation was subtracted from the 3 pre-activation frames for each cell.

#### Cell segmentation and spatial profile extraction of CRY2, myosin and NuMA fluorescence intensity

For spatial analysis of CRY2, myosin and NuMA fluorescence intensity along the cell periphery shown in (Fig5B, S6C, E), images were first corrected for photobleaching and registered to correct for XY drift as described above. Images were then segmented using a Fiji plugin-JFilament2D **[10].** Briefly, this method is based on stretching active contours that deform to detect bright elements in an image. Using the segmented contours, fluorescence intensity was extracted for each pixel around the cell periphery using a custom-built Fiji plugin. The average intensity profile of all pre-activation frames and the average intensity of 5 post-activation frames was used for subsequent spatial analysis of fluorescence. We defined the centre of the region of activation for each pole and cropped 225 pixels (corresponding to 90°) on either side of the centre thus giving profiles spanning −90° to +90° with 0 being the centre of the pole and ±90 being the equator. To obtain the spatial profiles of CRY2 and myosin, we first averaged the two polar pre-activation intensity profiles and the two polar post-activation intensity profiles to obtain a mean pre-activation and post-activation intensity profile respectively. We next calculated the average value of the mean pre-activation profile (F0) and finally divided the mean post-activation trace (F) by F0 to obtain the normalized fold change traces (F/F0) at the poles.

For the spatial profile of NuMA, we defined the position of the two poles and cropped 225 pixels (corresponding to 90°) on either side thus giving profiles spanning −90° to +90° with 0 being the pole. The two polar intensity profiles were averaged to give a mean polar intensity profile. To obtain the intensity of NuMA at the equator, we first measured the width of the metaphase DNA plate and computed a mean width across all cells (mean width = 48 pixels, ~3μm). The two equators were then defined and X/2 pixels were cropped on either side to give two equatorial profiles. The two profiles were averaged to give a mean equatorial profile. We next calculated the average value of the mean equatorial profile (F0) and finally divided the mean polar intensity trace (F) by F0 to obtain the normalized fold-change trace (F/F0) at the poles.

#### Determining the length of astral microtubules

The length of astral microtubules in FigS6B, was calculated from immunostaining images of α-tubulin. Images were first rotated such that the spindle was always perpendicular to the horizontal and a maximum projection of the middle 7 slices was used for analysis. Astral microtubule length *l_m_* was manually measured in Fiji as the distance between the centrosome and the tip of the astral microtubule. The length data was plotted as a histogram showing the mean and the standard deviation of the distribution.

#### Probability distribution histograms

The probability distribution histograms (Fig5F-H), were obtained by binning the spindle angular positions *Φ,* pre-activation (t=0), 14 minutes post-activation (t=14’) and at anaphase onset in bins of 10 degrees. Probability histograms were obtained by dividing the number of cells in each bin by the total number of cells.

#### Mean square deviation (MSD) analysis

The MSD analysis in FigS6D was done as follows. For the control condition, we obtained the mean squared deviation (MSD) curves of individual cells for delays (Δ*t*) up to 15 mins in steps of 1 min. At a given delay, for time *t*, we calculated the squared difference in positions at time *t*+Δ*t* and at time *t* and computed an average over all times.

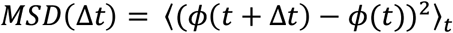

We then averaged the curves of individual cells and fitted the average MSD vs. delay curve to a saturating exponential function of the form 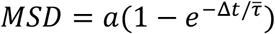 using nonlinear regression (*nlinfit* function in MATLAB) and estimated the values of *a* and 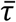. Here, *a* is the saturating value of the function and 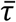 is the time constant in mins. Finally, we evaluated the diffusion coefficient (*D*) as the first derivative of the MSD at 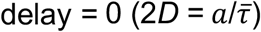.

#### Statistical and Data Analysis

All statistical analyses were performed using MATLAB_R2019a (Mathworks) or Excel. The Anderson-Darling test was used as a normality test to compare the distribution of the data sets. For normal distributions, we used unpaired Student’s *t*-test to test the difference between two independent datasets and paired Student’s *t*-test to test the difference between two dependent datasets. For non-normal distributions, we used Wilcoxon rank-sum to test the difference between two independent datasets and Wilcoxon sign-rank to test the difference between two dependent datasets. To analyze the differences among multiple experimental groups, we used one-way analysis of variance (ANOVA) test followed by post-hoc Tukey-Kramer test (for normal distributions) and Kruskal-Wallis test followed by post-hoc Dunn’s test (for non-normal distributions). To test differences between two distributions, 2-sample Kolmogorov-Smirnov test was used. Statistical significance was considered at *p*-values < 0.05 (*), <0.01 (**) and <0.001 (***). All boxplots show the median value (dark line), mean value (filled circle), the first and third quartile (bounding box) and the range (whiskers) of the distribution. Raw data points (light circles) are plotted on the side of all boxplots. Data are represented as Mean ± SEM except in FigS6E where it is represented as Mean ± SD. Number of cells analyzed for each experiment and the corresponding *p*-values are described in each figure legend.

## Supplementary figures and legends

**FigureS1.**
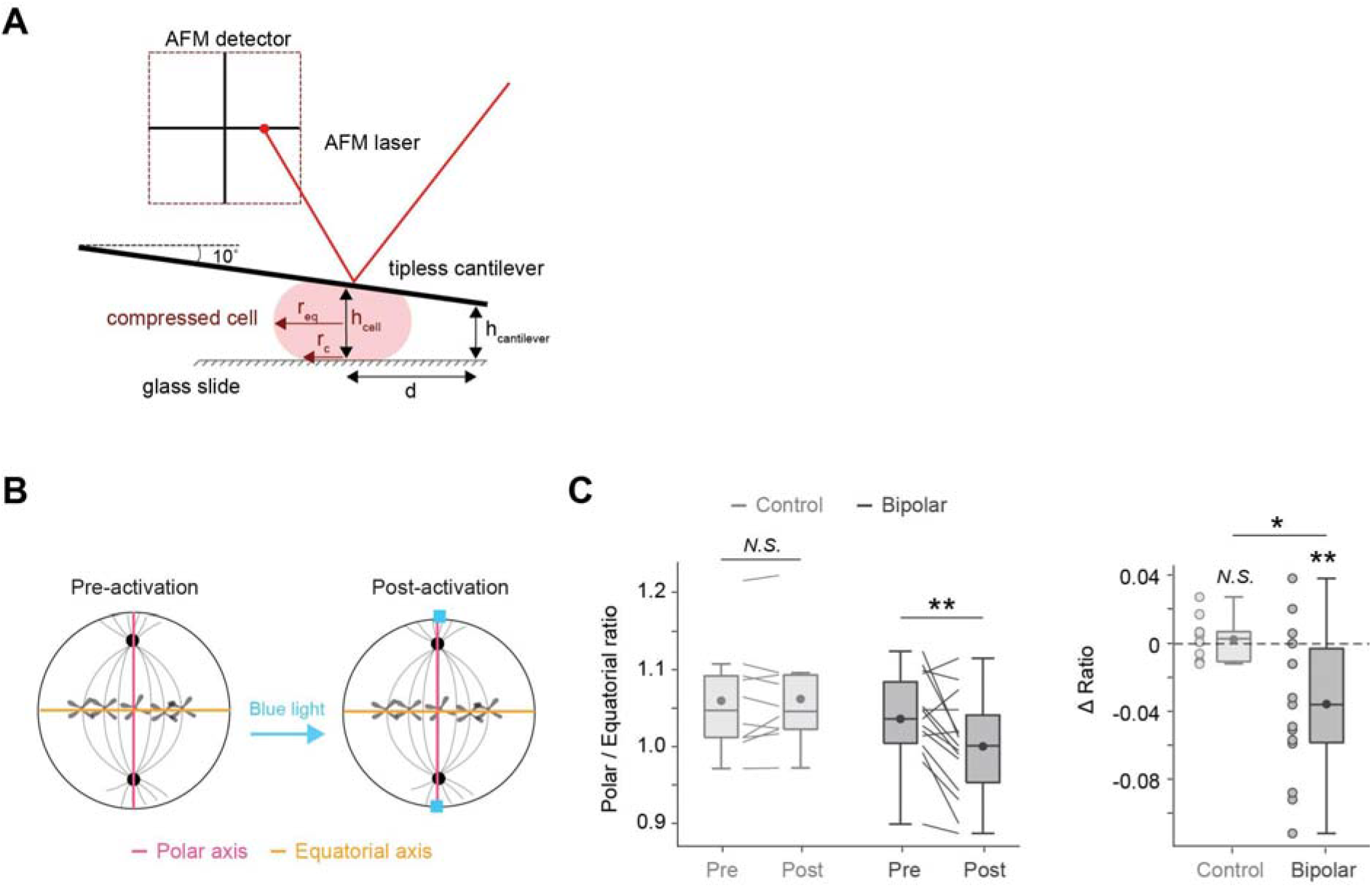
(Related to Figure1): Increase in RhoA activity increases cortical tension. A. Schematic depicting the AFM experiment. Single rounded MDCK cells were compressed under a tipless AFM cantilever. The cantilevers were mounted on a standard holder with a 10° tilt relative to the horizontal. The AFM laser measures the cantilever deflection which is converted into force knowing the cantilever’s spring constant. Confocal imaging is used to determine the cell radius at the midplane r_eq_ and the distance between the cantilever extremity and the cell centre *d*. Readings from the piezo height are used to determine the height of the extremity of the cantilever relative to the glass slide h_cantilever_ and the cell height h_cell_. See Methods for details. B. Schematic depicting the procedure used to measure shape change. We measured the length of the polar (pink) and equatorial (yellow) axes pre- and post-activation. We then compared the ratio of the length of polar over equatorial axis for cells pre- and post-bipolar optogenetic activation for cells as in Fig1J. C. Left: Boxplot showing the ratio of polar over equatorial axis pre- and 2 minutes post optogenetic activation in control and bipolar conditions. Paired students *t*-test, Control: *p*= 0.5911 (N.S), Bipolar: *p*= 0.0062 (**). Right: Boxplot showing the change in polar/equatorial axis of cells in response to optogenetic activation as in (C, Left). Student *t*-test compared to a 0% change in ratio: Control *p*= 0.5732 (N.S), Bipolar *p*=0.0061 (**); unpaired students *t*-test: control vs bipolar p= 0.0127 (*). Boxplots show median (dark line), interquartile range, mean (dark filled circle) and individual data points (light filled circle).

**FigureS2.**
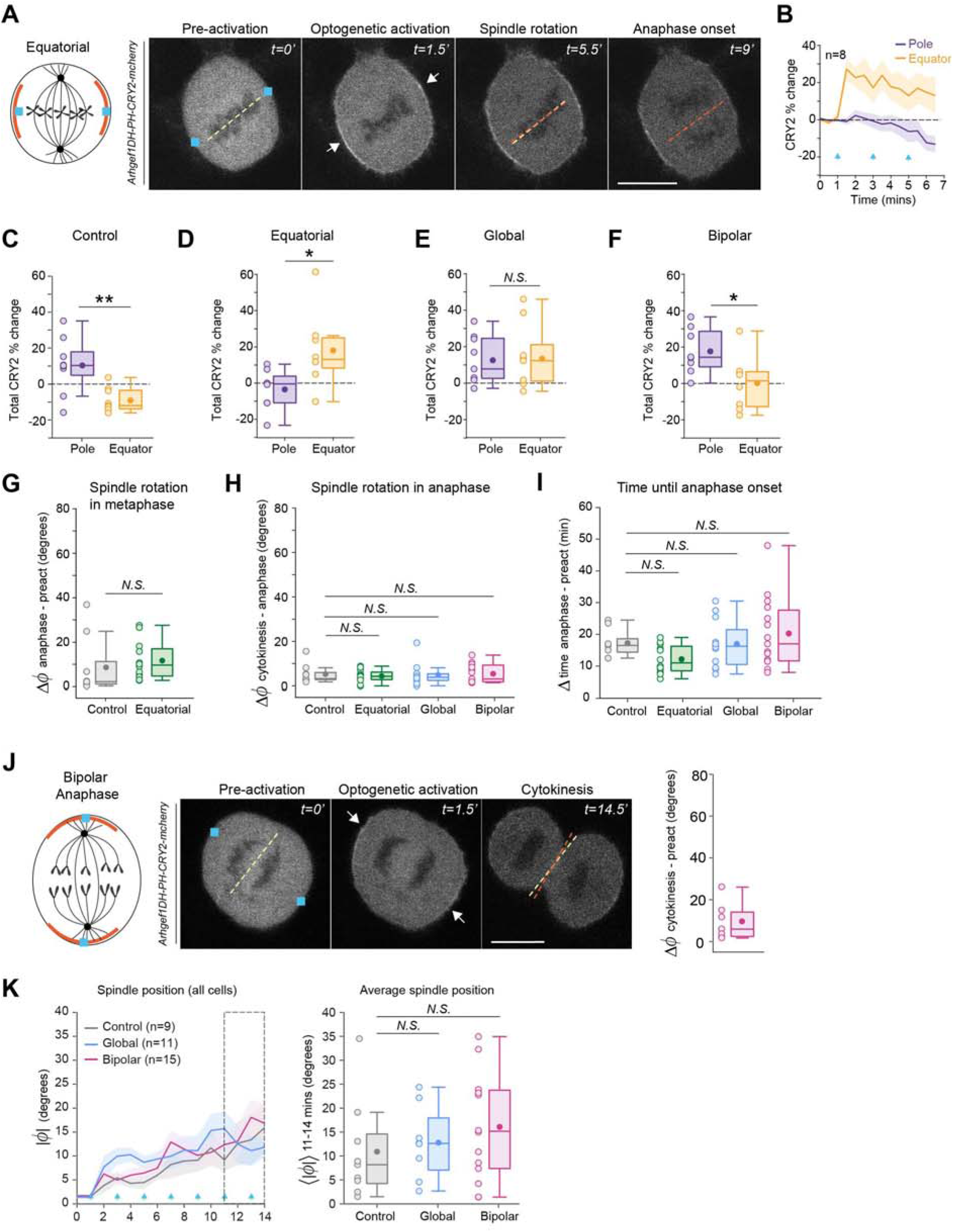
(Related to Figure1): Mitotic spindle orientation responds to inhomogeneities in RhoA activity and cortical tension. A. Left: Schematic showing the localisation of Arhgef1-DH-PH-CRY2-mCherry (red), after equatorial optogenetic activation in metaphase cells. Right: Time series showing localisation of GEF-DH-PH-CRY2-mCherry in cells before and after equatorial optogenetic activation until anaphase onset. Yellow and orange dashed lines indicate the position of the DNA in the metaphase plate before activation and at anaphase onset. Blue squares indicate the region of activation, white arrows point to regions of GEF accumulation. Scale bar = 10μm. Time is in minutes. B. Temporal plot showing the percentage change in GEF-DH-PH-CRY2-mCherry intensity at the poles and equator after equatorial activation. Plot shows mean ± SEM (n=8 cells). Blue arrows indicate the times of blue light stimulation. C. Boxplot showing the total percentage change in CRY2-mCherry intensity at the poles and equator under control condition. Boxplot shows median (dark line), interquartile range, mean (dark filled circle) and individual data points (light filled circle), (n=9 cells), student *t*-test, pole vs equator: *p*= 0.005 (**). D. Boxplot as in (C) after equatorial activation (n=8 cells), students *t*-test, pole vs equator: *p* = 0.029 (*). E. Boxplot as in (C) after global activation (n=9 cells), students *t*-test, pole vs equator: p= 0.833 (N.S). F. Boxplot as in (C) after bipolar activation (n=8 cells), students *t*-test, pole vs equator: *p* = 0.036 (*). G. Boxplot showing spindle rotation in metaphase for cells after control (n=9) and equatorial (n=12) activation. Boxplot shows median (dark line), interquartile range, mean (dark filled circle) and individual data points (light filled circle). Unpaired students *t*-test, Control vs Equatorial: *p*= 0.6629 (N.S). H. Boxplot showing spindle rotation in anaphase under control (n=9), equatorial (n=12), global (n=12) and bipolar (n=15) activation conditions for cells as in (1J and S2G). Boxplot shows median (dark line), interquartile range, mean (dark filled circle) and individual data points (light filled circle). Kruskal-Wallis and post-hoc Dunn’s test: Control vs Bipolar: *p*=1 (N.S); Control vs Equatorial: *p*= 0.9993 (N.S); Control vs Global: *p*=0.9928 (N.S). I. Boxplot showing duration of spindle rotation under control (n=9), equatorial (n=12), global (n=12) and bipolar (n=15) activation conditions for cells as in (S2H), computed as the time between initial activation and anaphase onset. Boxplot shows median (dark line), interquartile range, mean (dark filled circle) and individual data points (light filled circle). ANOVA and Tukey-Kramer test: Control vs Bipolar: *p*=0.8821 (N.S); Control vs Equatorial: *p*= 0.5771 (N.S); Control vs Global: *p*=1 (N.S). J. Left: Schematic showing the localisation of GEF-DH-PH-CRY2-mCherry (red) after bipolar optogenetic activation in early anaphase. Middle: Montage showing localisation of GEF-DH-PH-CRY2-mCherry in cells before and after bipolar optogenetic activation until cytokinesis. Yellow and orange dashed lines indicate the position of the DNA in the metaphase plate at the start of anaphase before activation and at the time of cytokinesis. Blue squares indicate the region of activation, white arrows point to regions of GEF-DH-PH accumulation. Scale bar = 10μm. Time is in minutes. Right: Boxplot showing spindle rotation in anaphase (n=7 cells). K. Left: Trajectories of absolute spindle angles before and 14 minutes after optogenetic activation for cells as in Fig1J. Right: Boxplot showing an average of the absolute spindle angle between 11-14 minutes. Control (n=9), bipolar (n=15), global (n=11). ANOVA and Tukey-Kramer test: Control vs Bipolar: *p*=0.46 (N.S); Control vs Global: *p*=0.92 (N.S).

**FigureS3.**
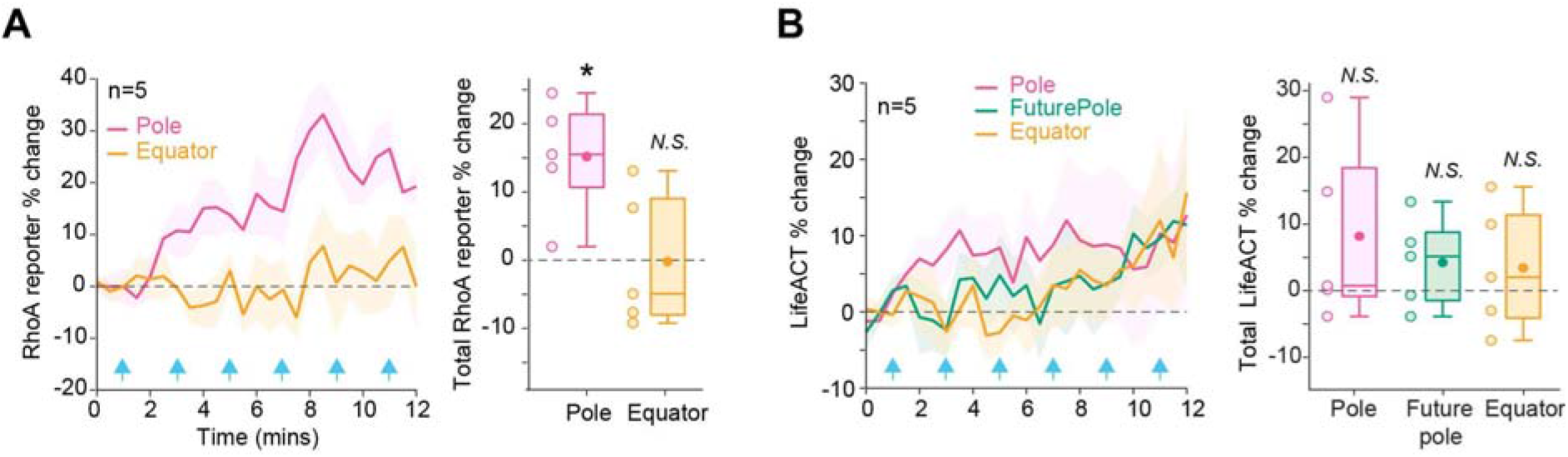
(Related to Figure 2): Mitotic spindles rotate away from myosin-enriched tensed cortical regions. A. Left: Temporal plot showing the percentage change in iRFP-RhoA biosensor intensity at the pole and equator after bipolar activation. Plot shows mean ± SEM (n=5 cells). Blue arrows indicate the times of blue light stimulation. Right: Boxplot showing the total percentage change in iRFP-RhoA biosensor intensity for cells as in (A, Left). Boxplot shows median (dark line), interquartile range, mean (dark filled circle) and individual data points (light filled circle). Student *t*-test compared to 0% change: Pole: *p*=0.0163 (*), Equator: *p*=0.9672 (N.S). B. Left: Temporal plot showing the percentage change in LifeAct-iRFP intensity at the pole, future pole and equator after bipolar activation. Blue arrows indicate the times of blue light stimulation. Plot shows mean ± SEM (n=5 cells). Right: Boxplot showing the total percentage change in LifeAct-iRFP intensity for cells as in (B, Left). Boxplot shows median (dark line), interquartile range, mean (dark filled circle) and individual data points (light filled circle). Student *t*-test compared to 0% change: Pole: *p*=0.2501 (N.S), Future pole: *p*=0.2329 (N.S), Equator *p*=0.4590 (N.S).

**FigureS4.**
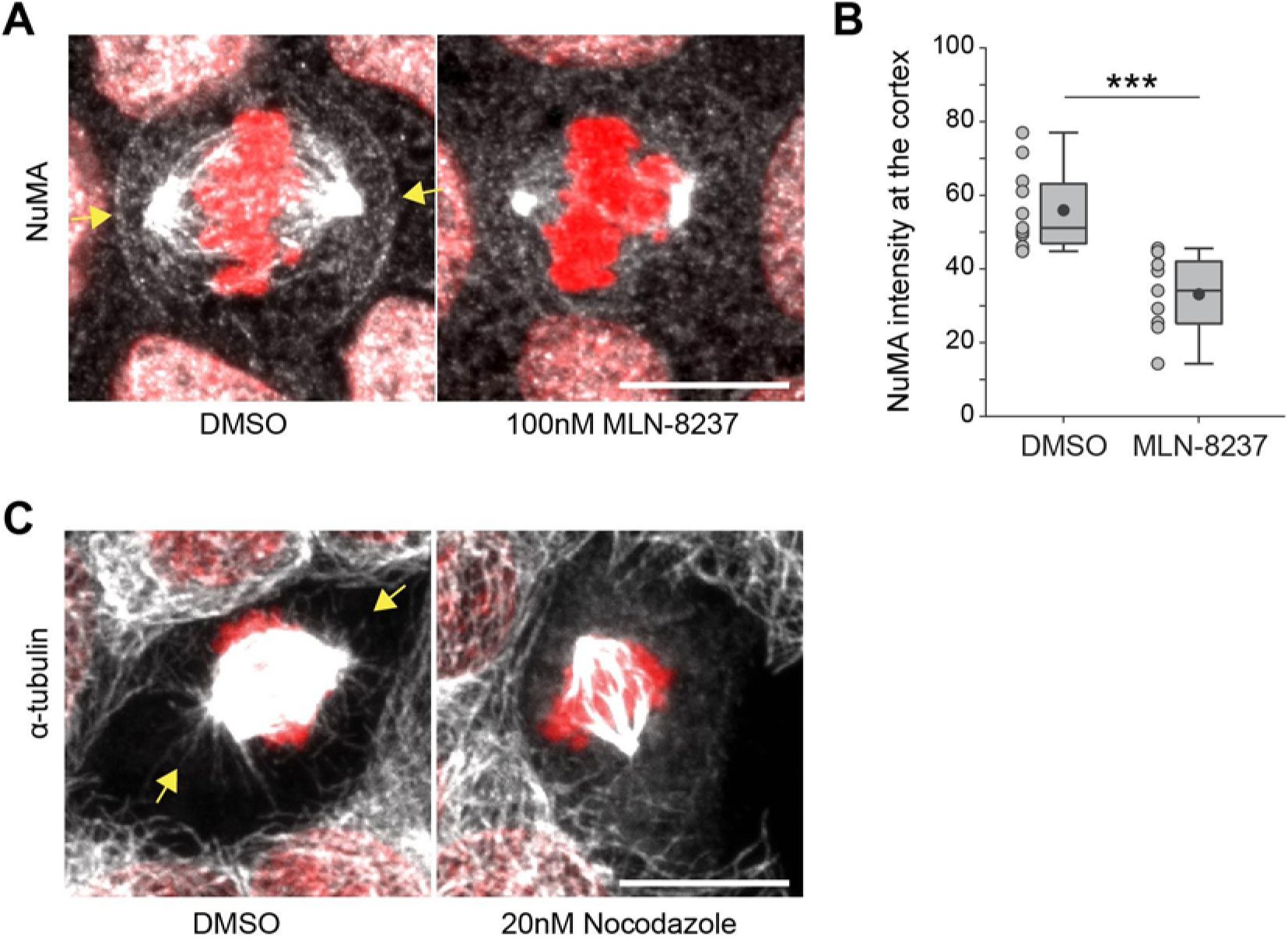
(Related to Figure3): Immunofluorescence images of NuMA and astral microtubules. A. Immunofluorescence images of WT MDCK cells stained for NuMA (greyscale) after treatment with DMSO and 100nM MLN-8237. Yellow arrows show cortical NuMA. DNA is stained with Hoechst (red). Scale bar =10μm. B. Cortical intensity of NuMA in cells treated with DMSO (n=11) and MLN-8237 (n=9) in arbitrary units. Boxplot shows median (dark line), interquartile range, mean (dark filled circle) and individual data points (light filled circle). Student *t*-test, DMSO vs MLN-8237: *p*= 0.000187 (***). C. Immunofluorescence images of WT MDCK cells stained for α-tubulin (greyscale) after treatment with DMSO and 20nM nocodazole to disrupt astral microtubules. Yellow arrows show the presence of astral microtubules. DNA is stained with Hoechst (red). Scale bar = 10μm.

**FigureS5.**
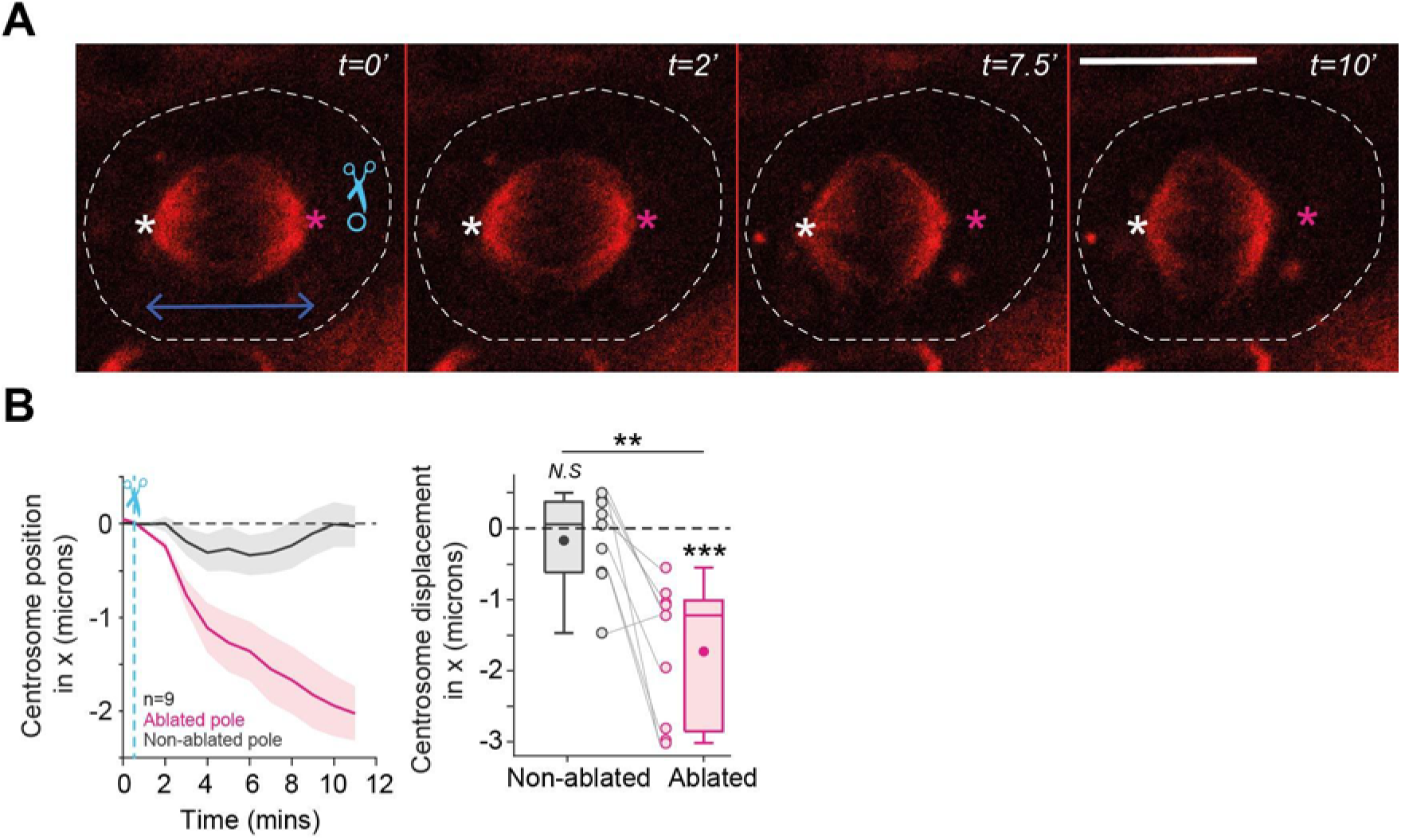
(Related to Figure 4): Centrosomes are subject to a net pulling force. A. Time series showing SiR-tubulin (red) in WT MDCK cells before and after laser ablation of astral microtubules on one side of the cell. Blue circle shows the ablated region. White and pink stars indicate the positions of the centrosomes at t=0’ on the non-ablated (white, NAB) and the ablated (pink, AB) side respectively. Scale bar = 10μm and time is in minutes. Experiment representative of n=9 cells. B. Left: Temporal plot showing the position of the two centrosomes along the x-axis following laser ablation for cells as in (A). The displacement of each centrosome is measured with the convention that movement towards the cell centre is negative and away from the centre is positive. Plot shows mean ± SEM (n=9 cells). Blue dotted line indicates the time of ablation. Right: Boxplot showing total centrosome displacement along the x-axis. Student *t*-test compared to 0 μm displacement: NAB *p*=0.4641 (N.S), AB p=7.3×10^-4^ (***); unpaired student *t*-test NAB vs AB *p*=0.0021 (**).

**FigureS6.**
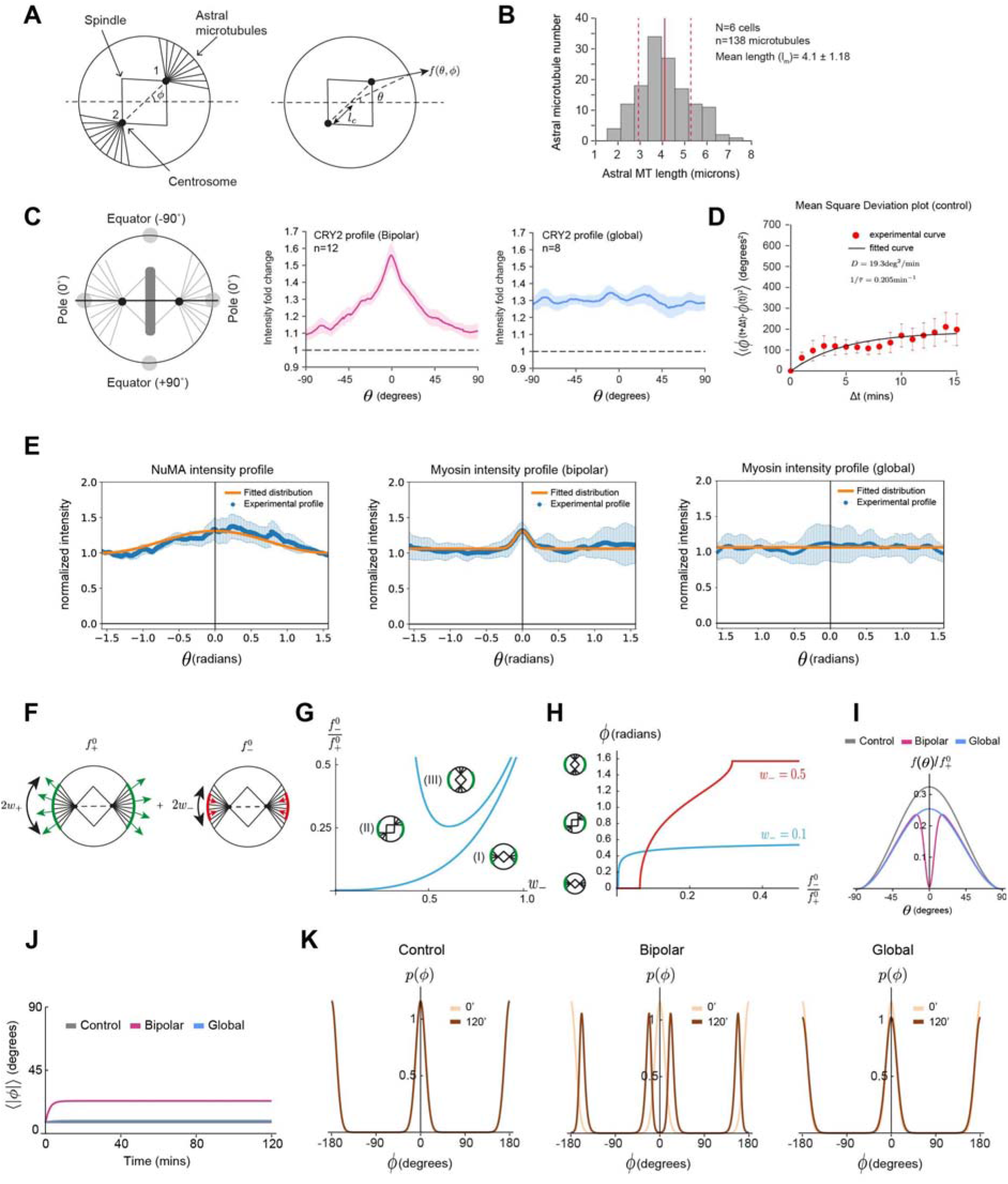
(Related to Figure5): Spindle rotation is an emergent property of patterned pulling forces exerted on astral microtubules. A. Schematics of the physical model of spindle rotation. The cell is assumed to have a circular shape. Left: Schematic of mechanical model for spindle rotation. The angle that the spindle makes with the horizontal is denoted *ϕ.* Astral microtubules emanate from the centrosomes (black dots) and reach the cortex when the centrosome-cortical distance is smaller than the maximal microtubule length *l_m_.* Right: A point on the cell cortex is denoted by its polar angle, *θ.* The distance from the cell center to a centrosome is denoted by *l_c_.* A microtubule contacting the cell cortex at angle *θ* is subjected to a force *f*(*θ, ϕ*). B. Histogram showing astral microtubule length distribution obtained from immunostaining images of NuMA. Solid red line indicates the mean length *l_m_* and dotted red lines indicate the mean± standard deviation. C. Left: Schematic of a cell with 0° representing the position of the pole and ±90° representing the position of the equator. Spatial profile of fluorescence intensity of CRY2 at the cortex post-activation normalised to pre-activation intensity after bipolar (n=12 cells, Middle) and global (n=8 cells, Right) optogenetic activation. Plot shows mean ± SEM. D. Mean square angle deviation (see methods for details) plotted as a function of time intervals for cells under control condition. Red circles show the experimental data, error bars show SEM and the solid black curve shows a fit to this data using Eq. 16 in Supplementary Theory. E. Fitting of experimentally measured normalized cortical profiles of NuMA and myosin after optogenetic activation for control, bipolar and global activation. Blue lines and error bars represent mean and standard deviations of experimental profiles, orange line: fitted distribution (see Eq. 27 - Supplementary Theory for details). F. Schematic of spindle subjected to patterned cortical forces. Here we assume that cortical forces acting on astral microtubules have a positive unperturbed contribution proportional to 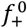, corresponding to pulling forces acting on microtubules (green arrows), and a negative perturbation proportional to 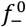, corresponding to reduced pulling forces acting on microtubules (red arrows). Forces act on each centrosome on a region with characteristic angles *w*_+_, *w*_. G. Phase diagram for stable configurations of the spindle, subjected to cortical forces as illustrated in (F). In region (I), the spindle angle *ϕ* = 0 is stable; in region (III), the spindle angle *ϕ* = *π*/2. is stable. In region (II), an intermediate configuration is stable. The width of the unperturbed pulling force region is taken from experimental measurements of cortical NuMA intensity profiles, *w*_+_ = 1.14 (Table 1-Supplementary Theory). H. Stable spindle angle as a function of the ratio 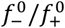, for two different values of *w*_. Other parameters as in (G). I. Normalized force profiles showing the ratio of 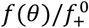 under control, bipolar and global activation conditions when the ratio 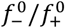 is set to 0.4. This ratio is chosen such that myosin recruitment post optogenetic activation results in a reduced pulling force without introducing a negative pushing force. J. Predicted average value of the spindle angle as a function of time under control, bipolar and global activation conditions, where the angle is measured between −90° and 90°, and the absolute value is averaged. K. Probability distribution *p*(*ϕ*) at t=0 and t=120 mins under control, bipolar and global activation.

**FigureS7.**
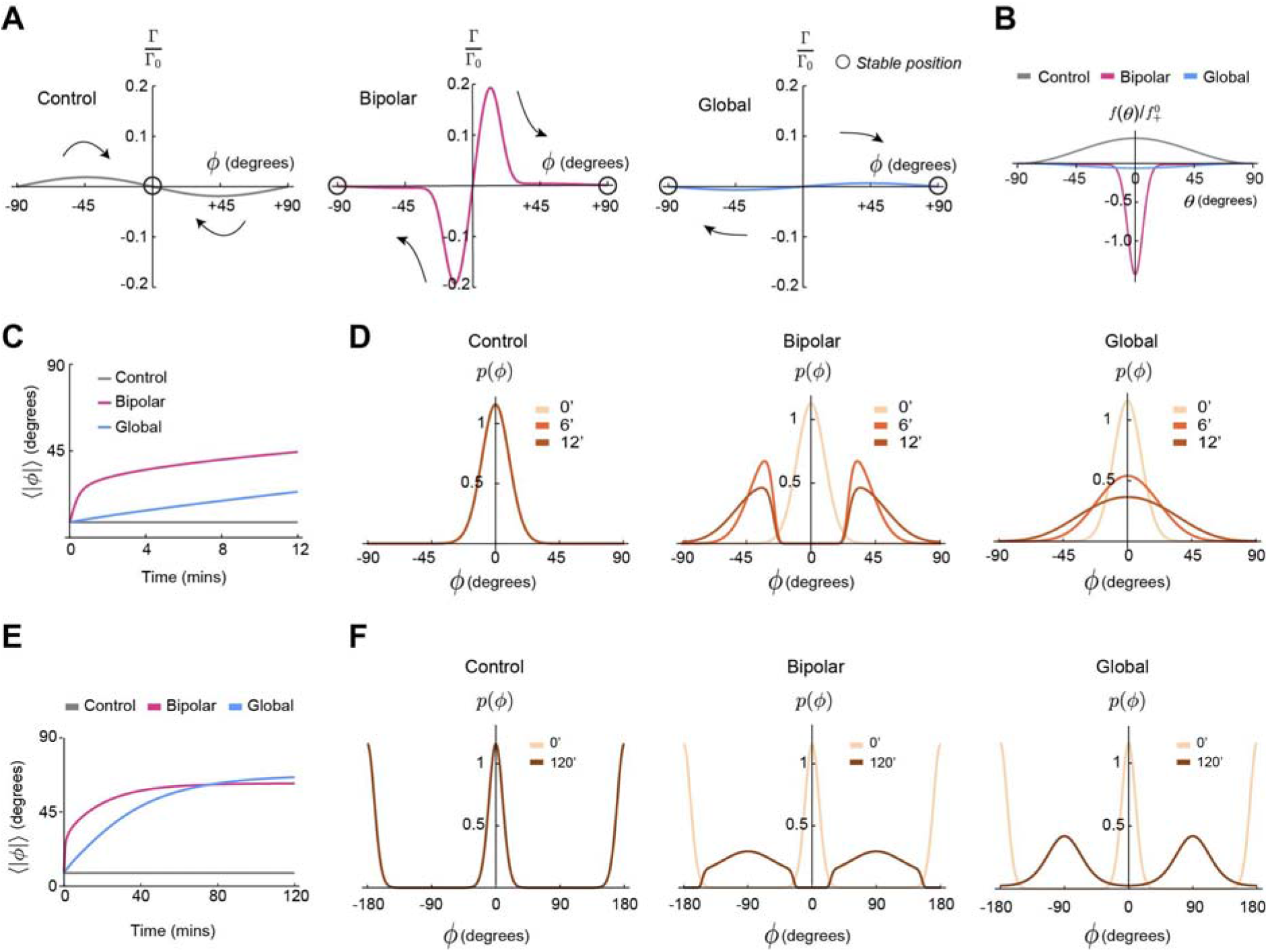
(Related to Figure5): A theoretical study of spindle response in a regime where pushing forces dominate. A. Predicted normalised torque (<u>Γ/Γ_o_</u>) as a function of spindle angle (*ϕ*) for control (Left), bipolar (Middle) and global (Right) activation. Arrows indicate the direction of spindle rotation. Stable spindle positions are indicated by black circles. Profiles of cortical pulling forces and reduced pulling forces are taken proportional to the fluorescence intensity profile of NuMA and myosin (B). In these simulations, 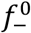 is chosen such that <u>*f*(*θ*)</u> is negative over the cell contour, representing a net pushing force exerted by astral microtubules on the cell cortex. See supplementary theory for further information. B. Normalized force profiles showing the ratio of 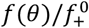 under control, bipolar and global activation conditions when the ratio 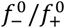 is set to 2.2. In this case, the force distribution *f*(*θ*) becomes negative under bipolar and global activation conditions corresponding to net cortical pushing forces. This parameter choice also results in larger rotation of the spindle. C. Predicted average value of the spindle orientation as a function of time after perturbation of the force profile for different conditions. The spindle orientation is measured between −90 ° and 90°, and the absolute value is averaged. D. Predicted probability distribution of the spindle angle, *p*(*ϕ*), at successive times (t=0,6,12 mins) after perturbation of the force profile under control, bipolar and global activation conditions. E. Predicted average value of the spindle angle as a function of time under control, bipolar and global activation conditions, where the angle is measured between −90° and 90°, and the absolute value is averaged. F. Probability distribution *p*(*ϕ*) at t=0 and t=120 mins under control, bipolar and global activation. in a regime where pushing forces dominate. At t=120 mins, the probability distribution is close to steady-state and has two broad peaks around 90° for both bipolar and global activation.

## SUPPLEMENTARY THEORY

In these notes, we describe a simple model for the rotation of a spindle which is subjected to patterned cortical forces, and its comparison to experiments. We introduce the model in section 1, discuss stable and unstable spindle position under a simple choice of patterned forces in section 2, and compare our results to spindle rotation experiments following optogenetic activation in section 3.

### 1. Model of spindle rotation in response to cortical forces

#### 1.1. Calculation of torque on the spindle

The 3D cartesian basis is denoted **e**_*x*_, **e**_*y*_, **e**_*z*_ and the spindle is assumed to move in the plane defined by **e**_*x*_, **e**_*y*_. We consider a spherical cell with radius *R* and use polar coordinates, such that a point on the cell contour is indicated by **r** = *R*cos *θ***e**_*x*_ + *R*sin *θ***e**_*y*_ (Fig. S6A). The spindle angle relative to the horizontal axis is denoted *ϕ*, and the distance from one centrosome to the cell center is denoted *l_c_*; such that the two centrosomes have positions:

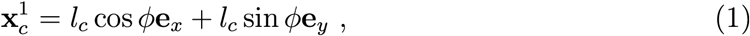

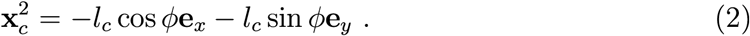

We denote **f**^1^(*θ,ϕ*), **f**^2^(*θ, ϕ*) the forces per microtubule, acting on an astral microtubule connected respectively to the centrosomes 1, 2 and touching the cell cortex at angle *θ* (Fig. S6A). We assume that this force is oriented along the microtubule connecting the centrosomes 1, 2 to a point on the cell surface. The force per microtubule emanating from centrosome 1, 2 can then be written:

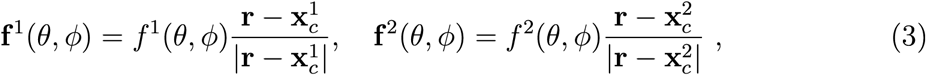

which defines the force magnitudes *f*^1^(*θ,ϕ*), *f*^2^(*θ,ϕ*). We consider here a situation where *f*2(*θ* + *π,ϕ* + *π*) = *f*^1^(*θ,ϕ*).

We denote *n*^1^(*θ*), *n*^2^(*θ*) the microtubule end density at a point *θ* on the cell contour, for microtubules emanating respectively from the centrosomes 1, 2. The angular microtubule density is taken to be uniform around the centrosome, with density *n_a_* = *N*_MT_/(2*π*) with *N*_MT_ the total number of microtubules around one centrosome. As a result the density of astral microtubules on the cell contour is

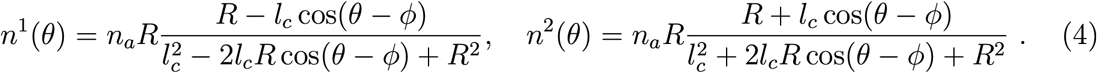

The total torque acting on the spindle is then given by **Γ** = Γ**e**_*z*_, with magnitude:

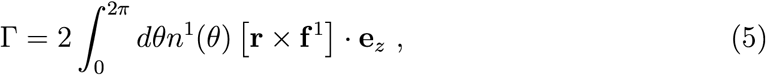

where the integral is taken over the microtubule density *n*^1^(*θ*) associated to centrosome 1, and the factor 2 arises from counting the effect of the two centrosomes, which here play symmetric roles. We then obtain the following explicit expression for the total torque:

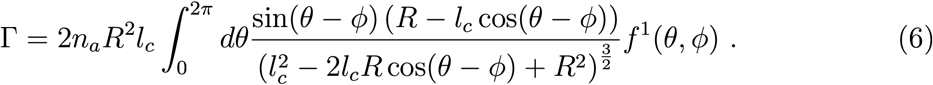

We note from the expression above that a uniform force distribution (*f*^1^(*θ, ϕ*) independent of *θ*) gives rise to a vanishing torque Γ = 0. Since we consider here a spherical cell, this is a consequence of invariance by rotation of the spindle, when the distribution *f* is not patterned on the cortex.

We now discuss the force distributions *f*^1^(*θ,ϕ*), *f*^2^(*θ,ϕ*). A force can be exerted only if the microtubules are long enough to reach the cortex from the centrosome. Denoting *l_m_* the microtubule length, which we take to be uniform, we then decompose the force distribution as:

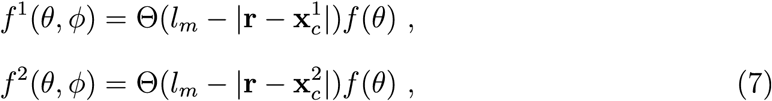

with Θ the Heaviside function, with Θ(*x*) = 0 for *x* < 0 and Θ(*x*) = 1 for *x* ≥ 0, and *f* (*θ* + *π*) = *f*(*θ*). In the right-hand side of Eq. 7, the first term ensures that only microtubules reaching the cortex are subjected to a force, and *f*(*θ*) is the force acting on a microtubule touching the cortex at angle *θ*, if the microtubule is long enough to come into contact with the cortex. The region where a force can be exerted on centrosome 1 is then within *ϕ* – *θ_m_* < *θ* < *ϕ* + *θ_m_*, with *θ_m_* defined by:

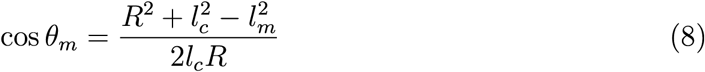

and the torque expression can be rewritten:

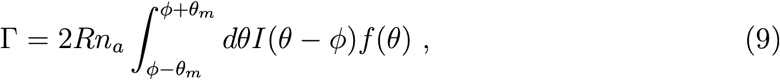

with 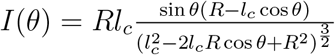 a dimensionless geometric function.

#### 1.2. Deterministic description of spindle rotation

One further assumes that the spindle rotation is subjected to dissipative forces, giving rise to an effective friction coefficient *γ*; such that spindle rotation is given by:

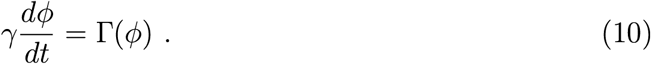

Steady-state spindle angles and their stability can be determined from Eq. 10.

#### 1.3. Stochastic description of the spindle rotation

We can alternatively consider the stochastic version of Eq. 10:

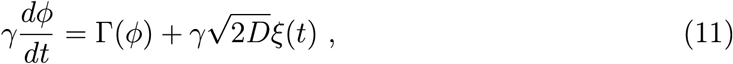

with *D* a diffusion coefficient, and *ξ*(*t*) a Gaussian white noise term satisfying

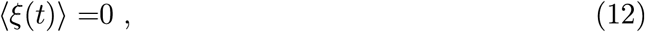

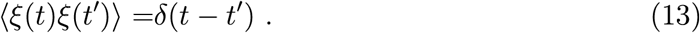

We now denote *p*(*ϕ*) the probability density function of observing the spindle angle *ϕ*. The corresponding Fokker-Planck equation for the evolution of *p*(*ϕ*) is [1]:

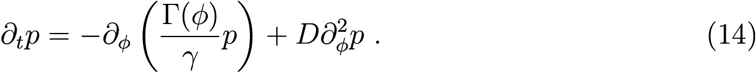

In a situation where *ϕ* = 0 is a stable steady-state, one can expand Γ(*ϕ*) for small deviations *ϕ* to obtain the Ornstein-Uhlenbeck equation:

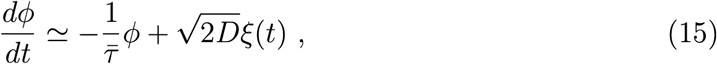

with 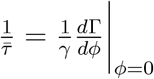. In that case, the mean-squared angle deviation reads for Δ*t* > 0, and assuming the process to be at stationary state:

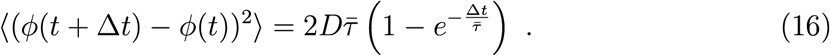

This relation can be used to extract values of 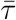 and *D* in control experiments from measurements of mean-squared spindle angle deviations (Fig. S6D).

### 2. Spindle orientation arising from a combination of patterned forces

Here we consider the case where the pattern of force distribution *f*(*θ*) arises from two contributions with different widths, according to (Fig. S6F):

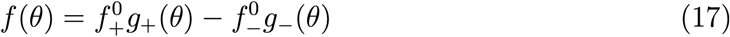

where *g*_+_(*θ*) and *g*___(*θ*) are two dimensionless functions, with normalization 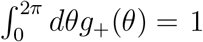 and 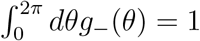. We think of the term in 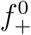 as the unperturbed pattern of cortical pulling forces associated with the distribution of NuMA at the cortex, and of the second term as a perturbation to this force distribution having a different spatial profile. This perturbation with magnitude 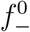 could arise from a contribution of cortical pushing forces with magnitude 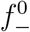 or from a local reduction of a pulling force pattern 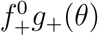.

We now discuss steady-state solutions of Eq. 10 and their stability. The total torque can be written:

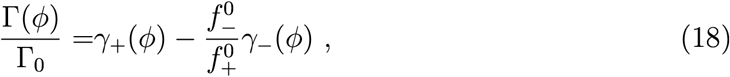

with 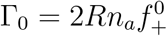 and

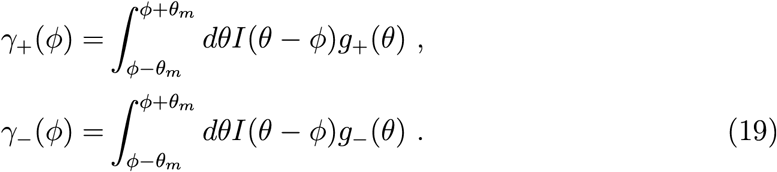

In practice we consider the functions *g*_+_ and *g*___ to be given by the periodic distribution:

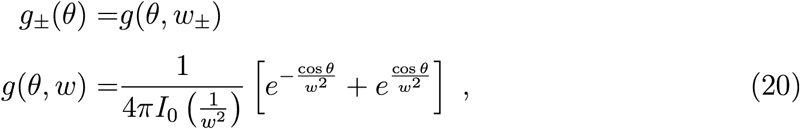

where *g*(*θ,w*) has maxima for *θ* = 0, *θ* = *π* (Fig. S6F) and verifies the normalization condition:

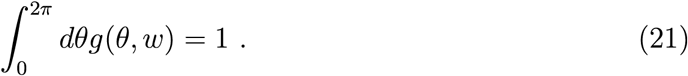

Following Eq. 10, a stationary solution for the spindle angle *ϕ* can be found by looking for solutions of the equation Γ(*ϕ*\*) = 0. The stability of the corresponding solution can be found by calculating the torque derivative with respect to the spindle angle *ϕ*:

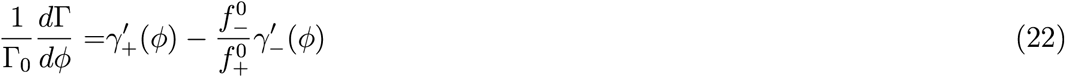

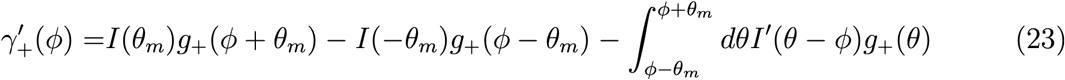

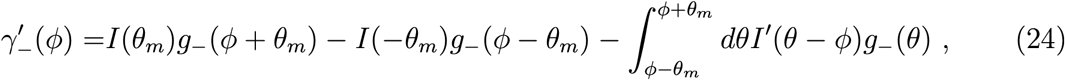

such that 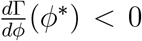 indicates a stable solution. With the choice of force distribution given in Eq. 20, the spindle angles *ϕ* = 0 and 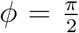 are always steady-state solutions, by symmetry. A change of stability of these solutions can be determined by solving for 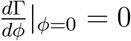 and 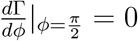, which using Eq. 22 corresponds to the conditions:

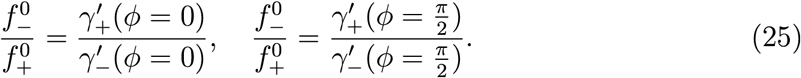

This relation is used to obtain curves in the phase diagram plotted in Fig. S6G which indicate the boundary of stability of *ϕ* = 0 and 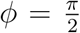. Here, the width of the pulling force region *w*_+_ is set to the experimentally measured width of the NuMA distribution (Table 1). In the region where both *ϕ* = 0 and 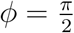 are unstable, an intermediate stable solution 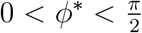 appears. Fig. S6H shows the stable solutions between 0 and *π*/2 as the magnitude of 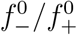 is increased for different widths of the perturbation profile *w*___. Overall we conclude that a pushing force pattern can lead to spindle rotation away from its original equilibrium point at *ϕ* = 0.

**Table 1.** Parameters fitted to experimentally measured NuMA and myosin cortical profiles (see Eq. 27). NA: non applicable.

| Profile | $a$ | $b$ | $w$ |
| --- | --- | --- | --- |
| NuMA | 0 | 7.24 | 1.14 |
| Myosin, bipolar | 1.06 | 0.12 | 0.1 |
| Myosin, global | 1.07 | NA | NA |

### 3. Comparison to experiments

To compare our model to experiments, we assume the following pattern of force distribution *f*(*θ*):

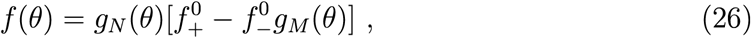

with *g_N_*(*θ*) corresponding to the profile of pulling forces, and the term in *g_M_*(*θ*) corresponds to a normalized reduction in cortical pulling forces or emergence of pushing forces, following optogenetic activation. Here we assume that local myosin recruitment after optogenetic activation results in a decrease in cortical pulling force, which is proportional to the NuMA concentration and to the myosin distribution. This is line with observations that treatment with MLN-8237 reduces recruitment of NuMA to the cell periphery and blocks rotation following optogenetic activation, suggesting that myosin recruitment in the absence of NuMA does not result in a force distribution driving spindle rotation (Figs. 3B, S4A, S4B).

We now discuss parameter values and comparison of theory predictions to experiments.

- The cell radius is taken equal to *R* = 8.9*μ*m, from measurements giving *R* = 8.9 ± 0.9*μ*m (mean±std, *N* = 51 cells).
- The distance from a centrosome to the cell center is taken equal to *l_c_* = 5.4*μ*m. This number is obtained from measurements of centrosome-centrosome distance in immunostained NuMA images, which gives a distance *2l_c_* = 10.3 ± 1.4*μ*m (mean±std, *N* =18 cells).
- The average astral microtubule length *l_m_* is determined from microtubule immunostaining images, which gives 4.1 ± 1.18*μ*m (n=138 microtubules, N=6 cells; Fig. S6B).
- We experimentally measured normalized NuMA intensity profiles and normalized myosin profiles (Fig. S6E) following optogenetic activation. NuMA profiles are normalized to the NuMA intensity at the cell equator, while myosin profiles after optogenetic activation are normalized to myosin profiles prior to optogenetic activation. Normalised NuMA and myosin intensity profiles are then fitted to the function:

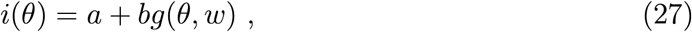

with *g*(*θ, w*) defined in Eq. 20; and *a, b, w* are free fitting parameters for the myosin profile in the bipolar activation condition, while we enforce *a* = 0 and leave *b* and *w* free parameters for the NuMA profile, and we enforce *b* = 0 for the myosin profile in the global activation condition, corresponding to a fit to constant profile. Corresponding parameters are reported in Table 1. This fitting procedure defines the NuMA intensity profile *i_N_*(*θ*) and the myosin intensity profiles *i*_*M*,bipolar_(*θ*) and *i*_*M*,global_(*θ*) for bipolar and global activation, respectively. The NuMA profile is then defined as

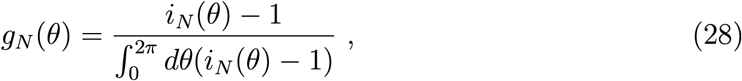

such that with this definition, we impose the normalisation 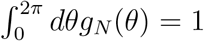; and we assume that the equatorial level of NuMA does not result in a pulling force. Indeed, to determine the distribution of NuMA stably anchored to the membrane and astral microtubules, we removed cytoplasmic background by permeabilisation-fixation and immunostained with an antibody against NuMA; in these images, NuMA shows a distinct peak at the poles but was indistinguishable from background at the equator (Fig. S4A). Therefore here we substract the equatorial level to NuMA fluorescence intensity profiles at the cortex. The perturbation profiles *g_M_*(*θ*) are defined as

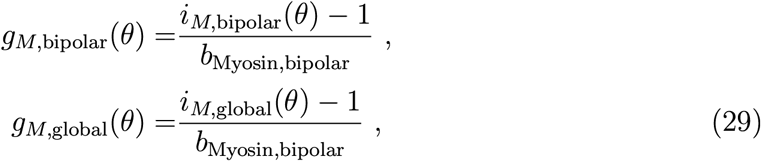

such that *i*_*M*,bipolar_(*θ*) = 1 or *i*_*M*,global_(*θ*) = 1, corresponding to a post-activation profile equal to the pre-activation profile, does not result in a perturbation of the cortical force distribution. The normalisation of *g*_*M*,bipolar_(*θ*) and *g*_*M*,global_ by *b*_Myosin,bipolar_ simply redefines the parameter 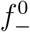.
- By comparing Eq. 16 to measurements of mean squared angle deviations in control experiments, we find *D* = 5.9 × 10^-3^ rad^2^.min^-1^ and 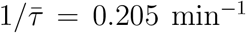 (Fig. S6D). In simulations described below we impose this measured value of *D* and use the measurements of 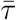 to set the value of *τ* = *γ*/Γ_0_. Indeed in the control case where the orientation *ϕ* = 0 is stable, we can use the approximation Eq. 15 and obtain

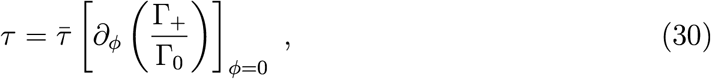

where Γ_+_ is the torque arising from unperturbed cortical forces, setting 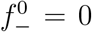 in Eq. 26. The ratio Γ_+_/Γ_0_ depends only on the geometrical parameters *R, l_c_, l_m_*, WuMA, allowing to relate *τ* to 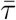; we find *τ* ≃ 0.2 min. The Fokker-Planck equation 14 can then be rewritten:

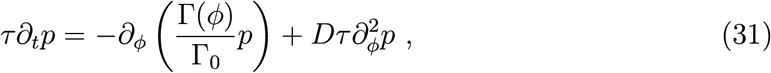

and can be solved with the knowledge of *τ, D*, geometrical parameters and force profiles determining Γ(*ϕ*)/Γ_0_, and using periodic boundary conditions on the domain −*π* < *ϕ* < *π*.

To obtain predicted torques for different conditions, we then combine Eq. 9, Eq. 26 and Eq. 29, and obtain:

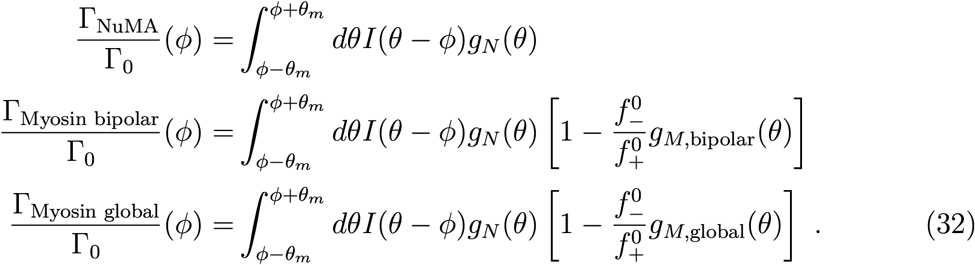

The ratio 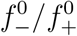 is a free parameter, which we set to 0.4 in Fig. 5C, where we plot the function Γ(*ϕ*)/Γ_0_ for the different conditions, as in Eq. 32. Here, this ratio has been chosen such that the perturbation introduced by myosin reduces the pulling force without introducing a negative, pushing force (Fig. S6I). We also show an example in which 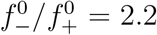 in Fig. S7. In that case, the force distribution *f*(*θ*) becomes negative after bipolar and global optogenetic activation, corresponding to a net pushing cortical force on microtubules (Fig. S7B). This parameter choice also results in a larger rotation of the spindle.

In Figs. 5E and S6K, S7D, S7F, we solve Eq. 14 for the probability distribution function *p*(*t, ϕ*) for different force distributions given in Eq. 32, with the initial condition *p*(*t* = *0,ϕ*) = *p*_0_(*ϕ*). *p*_0_(*ϕ*) is defined here as the steady-state distribution of *ϕ*, when the torque arises entirely from a force distribution given by the NuMA intensity profile.

In Figs. 5D and S6J, S7C, S7E, we plot the average value:

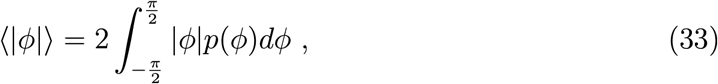

which would be obtained by measuring the spindle angle between −*π*/2 and *π*/2, taking the absolute value of the angle and averaging.

